# Interferon-Induced Transmembrane Protein 3 Blocks Fusion of Diverse Enveloped Viruses by Locally Altering Mechanical Properties of Cell Membranes

**DOI:** 10.1101/2020.06.25.171280

**Authors:** Xiangyang Guo, Jan Steinkühler, Mariana Marin, Xiang Li, Wuyuan Lu, Rumiana Dimova, Gregory B. Melikyan

**Author notes:** Corresponding author: Gregory B. Melikyan.

## Abstract

Interferon-induced transmembrane protein 3 (IFITM3) potently inhibits entry of diverse enveloped viruses by trapping the viral fusion at a hemifusion stage, but the underlying mechanism remains unclear. Here, we show that recombinant IFITM3 reconstituted into lipid vesicles induces negative membrane curvature and that this effect maps to its small amphipathic helix (AH). We demonstrate that AH: (i) partitions into lipid-disordered domains where IAV fusion occurs, (ii) induces negative membrane curvature, and (iii) increases lipid order and membrane stiffness. Additionally, replacing the IFITM3 AH with AH from an unrelated protein did not compromise its antiviral activity. These effects on membrane properties correlate with the fusion-inhibitory activity, as exogenous addition of AH to insulin-producing cells reduces glucose-stimulated insulin secretion. Our results thus imply that IFITM3 inhibits the transition from hemifusion to full fusion by imposing an unfavorable membrane curvature and increasing the order and stiffness of the cytoplasmic leaflet of endosomal membranes.

## Introduction

Infection by enveloped viruses proceeds through fusion of the viral membrane with the target cell membrane. Viral fusion, which is a critical step for establishing infection, is mediated by viral fusion proteins that are transmembrane glycoproteins protruding from the virus envelope. When activated by binding to cellular receptors and/or by acidic pH in endosomes, viral fusion proteins undergo extensive conformational changes (Harrison, 2008). As a result of this refolding, the two contacting membrane leaflets merge to form a hemifusion diaphragm (Chernomordik & Kozlov, 2003), which allows lipid mixing between the contacting leaflets, and ultimately culminates in full fusion through the formation of a fusion pore. Finally, enlargement of a fusion pore allows the release of the viral content into the cytosol and initiates infection.

Interferon-induced transmembrane proteins (IFITMs) are host factors that broadly and potently inhibit infection of diverse enveloped viruses, including the Influenza A virus (IAV), Dengue virus, Severe Acute Respiratory Syndrome-associated coronavirus (SARS-CoV), Respiratory Syncytial Virus (RSV) and Ebola virus (EBOV) (Brass et al., 2009; Diamond & Farzan, 2013). The IFITM family includes IFITM1 that localizes predominantly at the plasma membrane (Huang et al., 2011; Mudhasani et al., 2013), as well as IFITM2 and IFITM3, which contain an endocytic signal in their cytoplasmic N-terminal domain and are predominantly found in late endosomes and lysosomes (Amini-Bavil-Olyaee et al., 2013; Feeley et al., 2011). IFITM3 alone is responsible for the bulk of antiviral effects of interferon in cell culture (Brass et al., 2009) and has been shown to be important for restricting viral replication in cell culture and *in vivo* (Bailey, Huang, Kam, & Farzan, 2012; Everitt et al., 2013; Everitt et al., 2012; Y. H. Zhang et al., 2013).

While the importance of IFITM3 in host antiviral defenses *in vitro* and *in vivo* is well-documented, its mechanism of action is not clearly defined. IFITMs are type II transmembrane proteins with a cytoplasmic N-terminus, followed by a hydrophobic membrane-associated region and a C-terminal transmembrane domain (Bailey, Kondur, Huang, & Farzan, 2013; Ling et al., 2016). A number of residues scattered across the cytoplasmic region of IFITM3 have been shown to be essential for antiviral activity (John et al., 2013). The most popular view of the mechanism of IFITM3’s antiviral activity is that IFITM3 creates “tough membranes” that are not conducive to fusion, most likely by altering membrane curvature and fluidity (Chesarino et al., 2017; John et al., 2013; Li et al., 2013; Lin et al., 2013), but direct evidences and detailed mechanisms are lacking. Using single virus tracking in live cells, we have previously discovered that IFITM3 traps the IAV at a “dead-end” hemifusion stage that does not culminate in full fusion (Desai et al., 2014). In other words, IFITM3 does not restrict the lipid-mixing stage of viral fusion, but rather inhibits the formation of a fusion pore.

There is evidence suggesting that IFITMs may inhibit viral fusion *via* an indirect mechanism that involves recruitment or modulation of other host factors. First, IFITM3 has been reported to bind to and inhibit the function of vesicle-associated membrane protein-associated protein A (VAPA) (Amini-Bavil-Olyaee et al., 2013), which disrupts cholesterol transport from late endosomes and causes its accumulation in these compartments. However, high cholesterol content itself does not appear to inhibit virus-endosome fusion (Desai et al., 2014; Lin et al., 2013; Wrensch, Winkler, & Pohlmann, 2014; Wu et al., 2020). Second, it has been proposed that IFITM3 works by recruiting zinc metalloprotease ZMPSTE24 to endosomes and that this effector protein is responsible for virus restriction (Fu, Wang, Li, & Dorf, 2017). However, several lines of evidence support a direct mechanism of IFITM3-mediated virus restriction. We have employed single virus tracking in live cells expressing a functional IFITM3 protein tagged with autofluorescent proteins and shown that IAV enters IFITM3-enriched endosomes where it undergoes hemifusion, but fails to complete the fusion process (Suddala et al., 2019). We found that, by contrast, the IFITM3-resistant Lassa virus enters through a distinct pathway that utilizes endosomes devoid of IFITM3. Moreover, IFITM3 incorporation into the viral membrane inhibits fusion mediated by both IFITM3-sensitive and -resistant viral glycoproteins (Suddala et al., 2019; Tartour et al., 2017). These results suggest that IFITM3 inhibits viral fusion by a proximity-based mechanism that requires the presence of this factor at the sites of viral fusion.

Here, using liposome-based *in vitro* reconstitution assays, we provide evidence that IFITM3 restricts membrane fusion by inducing unfavorable negative curvature and stiffening the cytoplasmic leaflet of a target membrane at the site of viral hemifusion and that these effects map to the small amphipathic helix region of IFITM3.

## Results

### IFITM3 induces negative membrane curvature *in vitro*

To understand the molecular mechanism by which IFITM3 inhibits viral fusion, we sought to purify recombinantly expressed IFITM3 and its derivatives, reconstitute these into liposomes and assess their effects on the properties of lipid bilayers. For the purification purposes, we tagged IFITM3 with an N-terminal Strep-tag (Figure 1A) and verified its antiviral activity by expressing it in HEK 293T cells (Fig. S1A) and testing its ability to inhibit the influenza A virus (IAV) infection (Figure S1B). Next, the tagged IFITM3 was expressed in *Escherichia coli*, extracted by Triton X-100 and purified (Figure 1B). In order to visualize the IFITM3 association with membranes, we expressed and purified a fluorescently-tagged IFITM3 (referred to as IFITM3-iEGFP, Fig. 1A), with EGFP inserted into the N-terminal region of the protein (Suddala et al., 2019). We have previously shown that EGFP insertion does not compromise the antiviral activity of IFITM3 (Suddala et al., 2019). The purified proteins were then incorporated into preformed large unilamellar vesicles (LUVs) made of 16:0-18:1 phosphatidylcholine (POPC) and cholesterol, using a detergent-mediated reconstitution protocol (Rigaud & Levy, 2003). Here, detergent removal from a lipid/protein mixture through hydrophobic adsorption onto Bio-Beads SM-2 triggers protein insertion into lipid bilayers to generate proteoliposomes. Density gradient LUV flotation showed that both IFITM3 and IFITM3-iEGFP were efficiently incorporated into the liposomes (Figure S1C).

**Figure 1.**
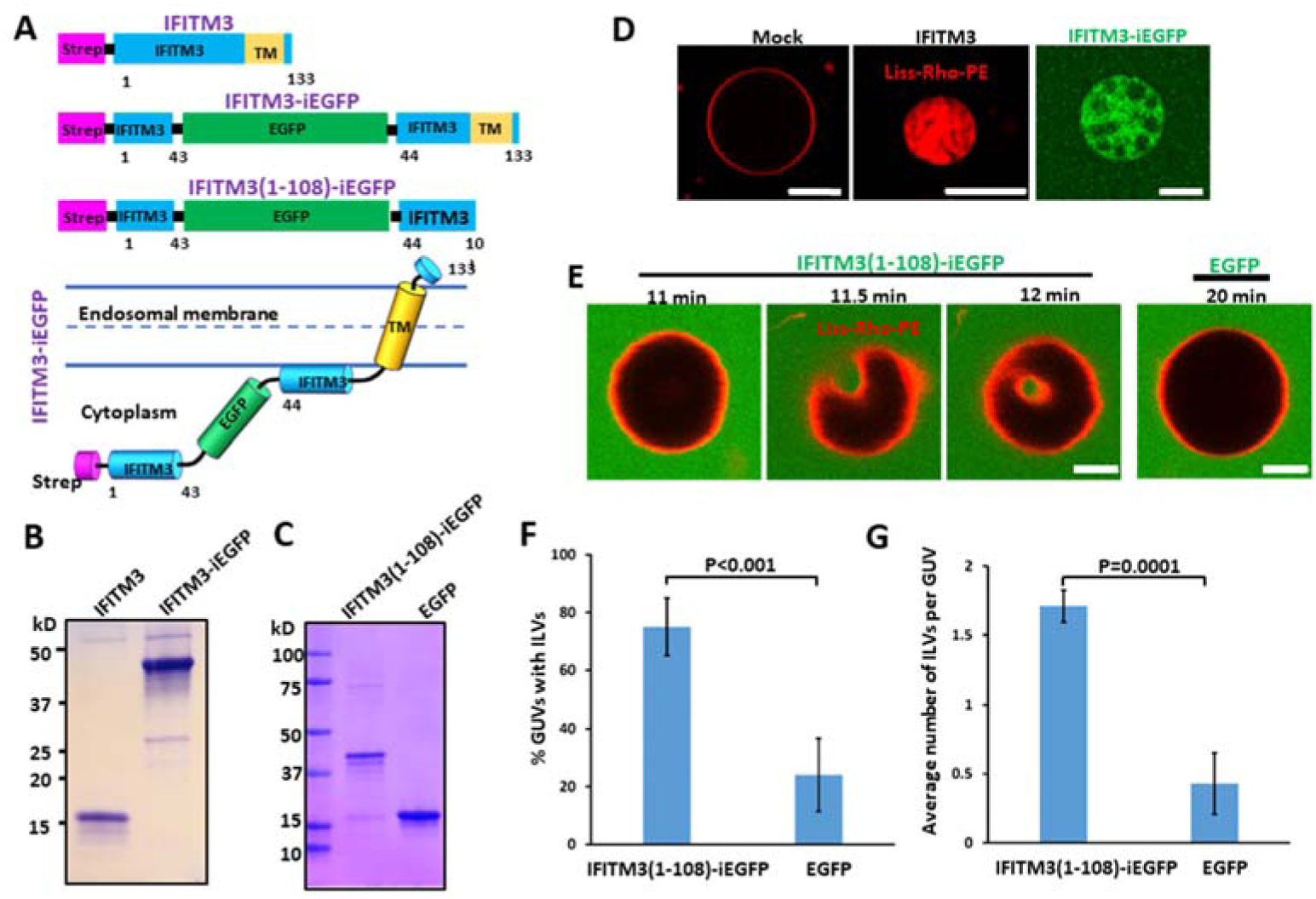
IFITM3 induces negative membrane curvature *in vitro*. (A) Illustration of domains of recombinant IFITM3 with an N-terminal Strep tag (IFITM3), Strep-tagged IFITM3 with an internal GFP tag (IFITM3-iEGFP), and C-terminally truncated IFITM3 lacking the transmembrane domain (TM) with Strep tag and internal GFP tags (IFITM3(1-108)-iEGFP). Lower cartoon illustrates the membrane topology of IFITM-iEGFP. The numbers indicate the amino acid numbers. (B, C) SDS–PAGE analysis and Coomassie blue staining of purified IFITM3 recombinant proteins shown in A. (D) IFITM3- and IFITM3-iEGFP-reconstituted LUVs (99.0 mol % POPC, 0.5 mol % cholesterol, 0.5 mol % Liss-Rho-PE) were dehydrated, electroformed into GUVs, and immediately imaged. Fluorescent Liss-Rho-PE lipid is shown in red. Scale bars 20 µm. (E) Time-lapse images of a GUV (99.0 mol % POPC, 0.5 mol % cholesterol, 0.5 mol % Liss-Rho-PE) incubated with 20 μM IFITM3(1-108)-iEGFP showing an inward budding of GUV membrane. A control GUV (right) was incubated with 20 μM EGFP and imaged under the same condition. Scale bars 5 µm. (F) Quantification of IFITM3-induced inward budding plotted as the percentage of GUVs with at least one IFITM3(1-108)-iEGFP or EGFP-filled intraluminal vesicle (ILV). Data represent mean ± SD from two independent experiments. At least 50 GUVs were analyzed per sample in each experiment. (G) Same as in (F), but the plotted values represent the average number of ILVs per GUV. See also Figure S1.

We next tested the sidedness of IFITM3 incorporation based upon its accessibility to proteolysis by trypsin. Addition of trypsin to IFITM3- or IFITM3-iEGFP-reconstituted liposomes resulted in a nearly complete cleavage of both molecules, similar to cleavage following proteoliposome solubilization in a detergent (Figure S1D). Thus, both reconstituted proteins appear to insert into liposomes in a preferential orientation, with their large cytoplasmic domains exposed to external medium. This result supports proper reconstitution of IFITM3 proteins in liposomes with an orientation similar to that in endosomal membranes (Bailey et al., 2013; Ling et al., 2016) (Fig. 1A).

In order to assess whether IFITM3 possesses membrane remodeling activity, IFITM3- or IFITM3-iEGFP-reconstituted proteoliposomes were dehydrated and electroformed into giant unilamellar vesicles (GUVs) using a previously described method (Girard et al., 2004). After formation, the N-terminally strep-tagged IFITM3 reconstituted into GUVs could be visualized by staining with the AlexaFlour-647 labeled streptavidin, further supporting its preferential orientation in lipid bilayers (Figure S1E). Importantly, we consistently observed that, in contrast to control GUVs, vesicles containing either IFITM3 or IFITM3-iEGFP protein contained intraluminal vesicles (ILVs) formed through inward budding of the GUV membrane (Figures 1D and S1E). The formation of ILVs or tubes from GUVs is a direct visualization of preferred or spontaneous membrane curvature (Karimi et al., 2018; Steinkuhler et al., 2020). This important finding implies that IFITM3 is capable of inducing negative membrane curvature, i.e. promoting membrane bending away from the protein (McMahon & Gallop, 2005).

We next asked whether the induction of negative membrane curvature by IFITM3 is mediated by its cytosolic domain (residues 1-108) (John et al., 2013; Ling et al., 2016). We therefore constructed a soluble IFITM3 fragment with GFP inserted into its N-terminal region (designated IFITM3(1-108)-iEGFP, Fig. 1A) and purified it without detergent solubilization (Figure 1C). Interestingly, IFITM3(1-108)-iEGFP showed very weak membrane binding to LUVs (<5% of protein input), as assessed by a liposome co-sedimentation assay (Julkowska, Rankenberg, & Testerink, 2013) (Figure S1F). Next, purified IFITM3(1-108)-iEGFP was added to preformed GUVs containing POPC and cholesterol prepared using a standard electroformation method (Angelova & Dimitrov, 1986). IFITM3(1-108)-iEGFP but not EGFP protein induced inward budding and formation of ILVs within 20 minutes of addition to GUVs, as illustrated by time-resolved imaging (Fig. 1E). As expected, intraluminal vesicles trapped IFITM3(1-108)-iEGFP from external solution (Figure 1E). Quantification of the inward budding effect of this protein showed that 75% of GUVs treated with IFITM3(1-108)-iEGFP contained ILVs and each GUV contained, on average, 1.7 ILVs (Figure 1F, G). By contrast, EGFP-treated GUVs rarely contained ILVs and any ILVs detected were not filled with EGFP, suggesting that these structures formed during GUVs electroformation. These results imply that the IFITM3 cytosolic domain induces negative membrane curvature, in spite of poor binding to membranes, and that the transmembrane domain is dispensable for this activity.

### IFITM3 amphipathic helix is responsible for induction of negative membrane curvature

It has been shown that a highly conserved small region (residues 59-68) of the IFITM3’s cytoplasmic domain, predicted to form an amphipathic helix (AH), is essential for antiviral activity (Chesarino et al., 2017; Z. Zhang, Liu, Li, Yang, & Zhang, 2012). Since many amphipathic helices can sense membrane curvature or induce membrane remodeling (Drin & Antonny, 2010), we asked whether the AH of IFITM3 is responsible for its ability to induce negative membrane curvature. To answer this question, we made a triple alanine substitutions S61A, N64A, T65A (designated IFITM3-3M, Figure 2A) in the AH region, which have been shown to greatly reduce the amphipathic moment of the AH and abrogate the antiviral activity against IAV (Fig. 2A) (Chesarino et al., 2017). The IFITM3-3M mutant was purified (Figure S2A) and reconstituted into GUVs (Figure 2B). We found that the inward budding activity of the GUV-reconstituted IFITM3-3M was significantly impaired compared to wild-type IFITMs. Only 7.1% of IFITM3-3M-reconstituted GUVs contained at least one ILV, as compared 81.7% of wild-type IFITM3-reconstituted GUVs (Figure 2B, C). We also tested the triple alanine mutant in the context of the IFITM3 cytosolic domain tagged with EGFP (IFITM3(1-108)-3M-iEGFP) and found that this soluble construct also exhibited impaired ILV-forming activity when added to preformed GUVs (Figure S2B-E). The above results map the negative curvature-promoting activity of IFITM3 to its AH.

**Figure 2.**
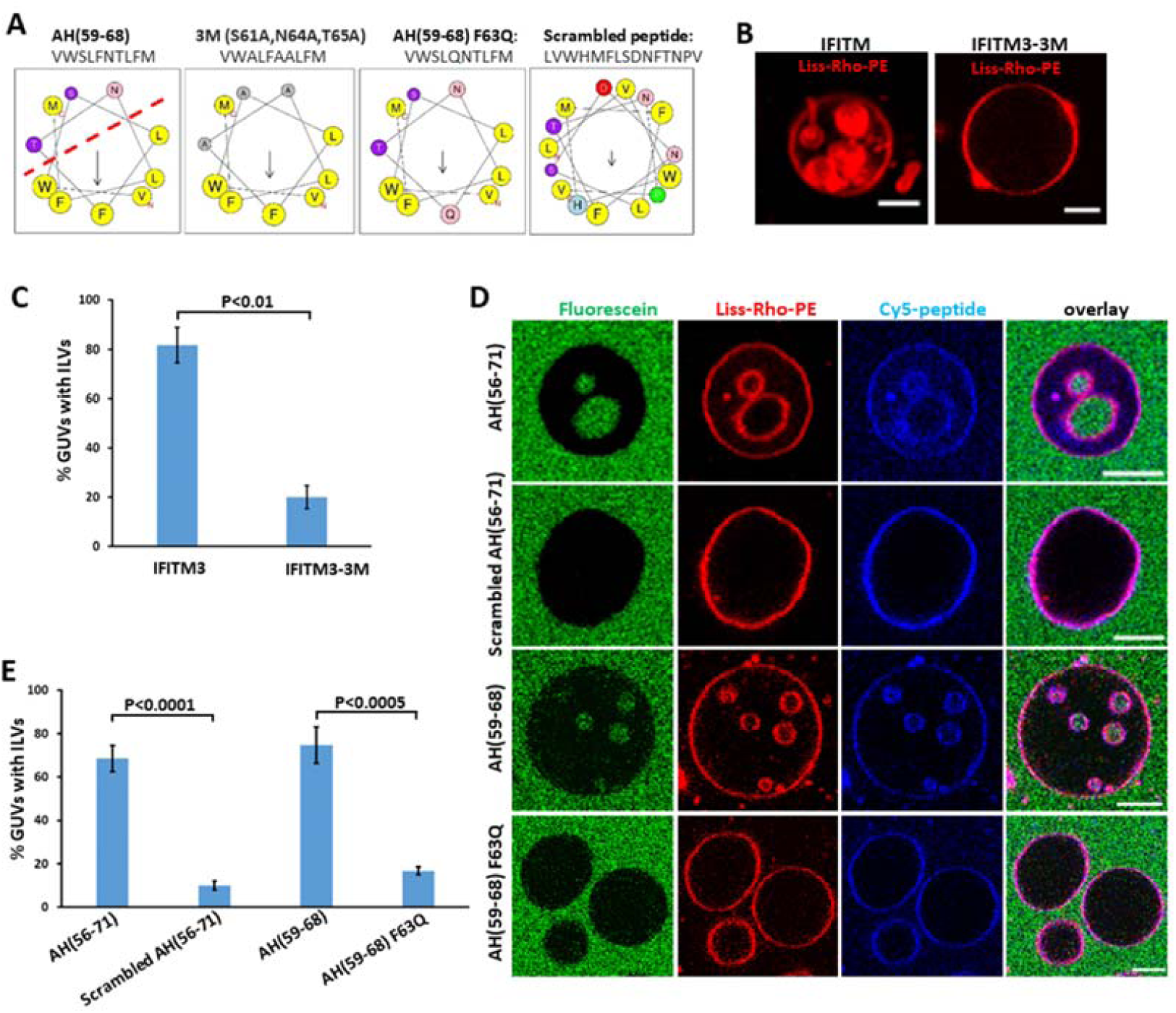
IFITM3 amphipathic helix is responsible for induction of negative membrane curvature. (A) Sequences and helical wheel projection plots created for IFITM3 AH and mutants using HELIQUEST software. Hydrophobic residues are displayed as gray or yellow, while hydrophilic residues are displayed as pink or purple. Arrows represent the magnitude and orientation of the mean hydrophobic moment value calculated by HELIQUEST software. (B) IFITM3- and IFITM3-3M-reconstituted LUVs were dehydrated, electroformed into GUVs and immediately imaged. Scale bars 10 µ m. (C) The percentage of GUVs, prepared and treated as in (B), containing at least one ILV. Data represent mean ± SD from two independent experiments, with at least 30 GUVs analyzed per sample in each experiment. (D) GUVs were treated with 10 µ M AH (56-71), 10 µ M Scrambled AH (56-71), 10 µM AH(59-68) or 30 µM AH(59-68) F63Q for 30 min and imaged. 0.3 µ M fluorescein was added to the external buffer to mark *bona fide* inward budding of GUVs. Scale bars 10 µ m. (E) Percentage of GUVs, prepared and treated as in (D), with at least one ILV containing fluorescein. Data represent mean ± SD of the results of three independent experiments, with at least 90 GUVs counted per sample in each experiment. See also Figure S2.

To determine whether IFITM3 AH alone is sufficient to induce negative membrane curvature, a peptide corresponding to this region (residues 59-68) and a longer peptide (residues 56-71) were synthesized and labeled with Cy-5 dye at their N terminus (Figure 2A). A peptide with scrambled amino acid sequence (Scrambled AH (56-71)) and the F63Q AH(59-68) mutant peptide (AH(59-68) F63Q) were used as controls.

Substitution of the hydrophobic F63 residue to glutamine has been shown to markedly decrease the peptide’s amphipathicity (Fig. 2A) and nearly abrogate its antiviral activity (Chesarino et al., 2017). The partition coefficients of the peptides, which reflect their membrane binding affinity, were measured using a liposome co-sedimentation assay (Figure S2F). The AH(56-71), AH(59-68) and Scrambled AH(56-71) peptides exhibited similar membrane binding affinities, with more than 80% of the input peptide binding to liposomes (at input peptide to lipid ratio 1:50). The AH(59-68) F63Q exhibited a slightly weaker membrane binding affinity, with 58.4% of the input peptide binding to liposomes.

Next, membrane binding and remodeling activity of these peptides added to GUVs were tested by imaging. All peptides were clearly enriched at the GUV membrane (Figure 2D), consistent with the membrane partitioning data (Fig. S2F). Strikingly, addition of AH(56-71) or AH(59-68) peptides to preformed GUVs containing POPC and cholesterol led to efficient inward budding, resulting in 68.5% and 74.7% of the GUVs containing at least one ILV, respectively (Figure 2D, E). These inner vesicles contained the aqueous marker fluorescein, which was added to GUVs externally, along with the peptides, in order to ensure that ILVs were formed *de novo* through inward budding (Figure 2D). Importantly, addition of the same concentration of the Scrambled AH(56-71) peptide or a 2-fold excess of the AH(59-68) F63Q peptide (to compensate for its slightly lower membrane affinity) did not induce considerable GUV vesiculation, with respectively 9.9% and 16.6% of the GUVs containing ILVs (Figure 2D, E). ILVs seemed to fission from the outer membrane as judged by their position in the GUVs, which is consistent with a recent finding of membrane fission of buds due to high membrane spontaneous curvature (Steinkuhler et al., 2020). Taken together, these results confirmed that IFITM3 AH alone is sufficient to induce negative membrane curvature and that this effect is critically dependent on the amphipathicity of this region.

### Negative membrane curvature induced by IFITM3 is facilitated by cholesterol and counteracted by lyso-lipids

Different lipids have different effective shapes manifested in spontaneous curvature that can impose positive or negative curvature to a lipid bilayer (Helfrich, 1973). Thus, biological membrane curvature is determined by an interplay between membrane proteins and lipids (Bassereau et al., 2018; Stachowiak, Hayden, & Sasaki, 2010; Steinkuhler et al., 2020). Lipids are categorized as “cylindrical”, “cone”, and “inverted cone” shaped. Cylindrical lipids, such as phosphatidylcholine (PC), prefer planar membranes, whereas cone-shaped lipids, such as cholesterol and phosphatidylethanolamine (PE), prefer negative membrane curvature, and inverted cone-shaped lipids, like Lyso-PC (LPC), promote positive membrane curvature (Kozlov, 2007).

We therefore tested whether lipids play a role in IFITM3-mediated GUV vesiculation. First, we electroformed POPC GUVs with or without cholesterol (0.5 mol%) and treated these with IFITM3(1-108)-iEGFP. Whereas this treatment resulted in 75.5% of cholesterol-containing GUVs with ILVs, a significant impairment of inward budding (8.5% GUVs with ILVs) was observed in the absence of cholesterol (Figure 3A, B). Similarly, addition of the AH(56-71) peptide to GUVs without cholesterol also failed to induce efficient inward budding (Figure S3A, B). Notably, IFITM3-induced inward budding in GUVs was independent of cholesterol concentration up to 20 mol% (Figure S3C). Importantly, AH(56-71) mediated inward budding of GUVs without cholesterol could be rescued by inclusion of another conical lipid, DOPE (20 mol%) (Figure S3D). This finding indicates that IFITM3-induced negative membrane curvature is facilitated by cone shaped lipids.

**Figure 3.**
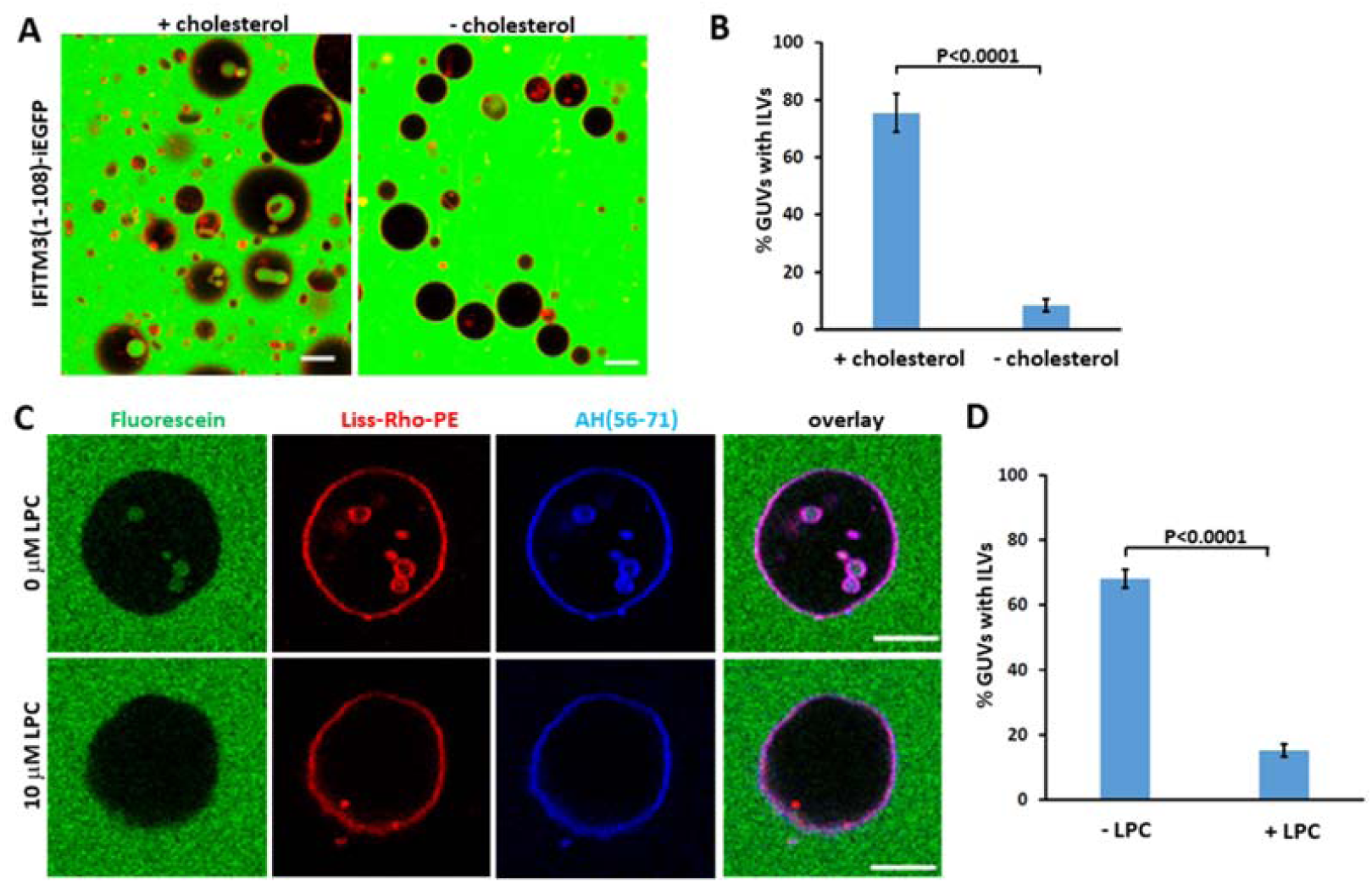
Negative membrane curvature induced by IFITM3 is facilitated by cholesterol and counteracted by lyso-lipids. (A) GUVs with cholesterol (99.0 mol % POPC, 0.5 mol % cholesterol, 0.5 mol % Liss-Rho-PE) or GUVs without cholesterol (99.5 mol % POPC, 0.5 mol % Liss-Rho-PE) were incubated with 20 μM IFITM3(1-108)-iEGFP for 30 min and imaged. Scale bars 20 μm. (B) Quantification of inward budding shows the percentage of GUVs prepared and treated as in (A) with at least one EGFP-positive intraluminal vesicle (ILV). Data represent mean ± SD of the results of three independent experiments, with at least 90 GUVs analyzed per sample in each experiment. (C) GUVs (99.0 mol % POPC, 0.5 mol % cholesterol, 0.5 mol % Liss-Rho-PE) were treated with 10 μM AH(56-71) and 10 μM LPC dissolved in methanol (+LPC) or the same volume of methanol (-LPC) for 30 min and imaged. Fluorescein (0.3 μM) was added to mark inward budding of GUVs. Scale bars 10 μm. (D) Quantification of inward budding showing the percentage of GUVs prepared and treated as in (C), with at least one fluorescein-containing intraluminal vesicle (ILV). Data represent mean ± SD of three independent experiments, with at least 35 GUVs analyzed per sample in each experiment. See also Figure S3.

We next tested the effect of a conical lipid, LPC, on GUV budding mediated by IFITM3. LPC was first titrated by adding to preformed GUVs containing POPC and cholesterol. As expected based upon positive curvature induced by this lipid (Fuller & Rand, 2001), we observed outward budding of the GUV membrane in an LPC dose-dependent manner (Figure S3E). A moderate concentration of LPC (10 µM) was then added together with the AH(56-71) peptide to preformed GUVs containing POPC and cholesterol. LPC abolished the AH(56-71)-induced GUV inward budding <u>(Figure 3C,</u> <u>D)</u>, showing that negative membrane curvature caused by IFITM3 is counteracted by the presence of conical lipid in the membrane.

### IFITM3 partitions into liquid-disordered membrane domains that support IAV fusion

Cell membranes contain lipid rafts, which are liquid-ordered domains with more tightly packed lipids than the non-raft, liquid-disordered phase of the bilayer (Rajendran & Simons, 2005a). Lipid rafts form a platform for signaling proteins and receptors and have been suggested to serve as potential virus entry sites (Chazal & Gerlier, 2003), but a formal proof is still lacking due to the fact that small and dynamic lipid microdomains are difficult to visualize in living cells. Post-translational modifications, such as S-palmitoylation, target transmembrane proteins to lipid rafts (Levental, Lingwood, Grzybek, Coskun, & Simons, 2010). Palmitoylation of the conserved cysteine residues downstream of the IFITM3 AH region regulate the membrane domain targeting and antiviral activity of this protein (Yount et al., 2010). We therefore asked whether IFITM3 has any preference to lipid microdomains and whether lipid rafts may play a role in IFITM3’s antiviral activity. To address this question, we utilized phase-separated GUVs as a model for microdomains within the endosomal membrane (Kaiser et al., 2009). GUVs containing sphingomyelin (SM), cholesterol and poly-unsaturated phosphatidylcholine (DOPC) segregate into lipid ordered (Lo) and lipid disordered (Ld) phases (Wesolowska, Michalak, Maniewska, & Hendrich, 2009) corresponding to lipid raft and non-raft microdomains of biological membranes, respectively. Incorporation of fluorescent markers partitioning into Lo (Top-cholesterol) or Ld (Liss-Rho-PE) phases allowed the visualization of the two phases (Figure 4A, B). The AH (56-71) peptide added to phase-separated GUVs partitioned to the Ld phase and caused inward budding from this domain, as evidenced by ILVs containing exclusively the Ld marker Liss-Rho-PE (Figure 4C). The Scrambled AH (56-71) peptide also exhibited an Ld phase preference, but failed to cause inward budding from this phase (Figure 4C). Intriguingly, palmitoylation of Cys71 of a fluorescently-tagged AH (56-75) peptide did not target this peptide to Lo phase. This peptide clearly partitioned to the Ld phase and appeared to concentrate at the phase boundary (Figure S4A).

**Figure 4.**
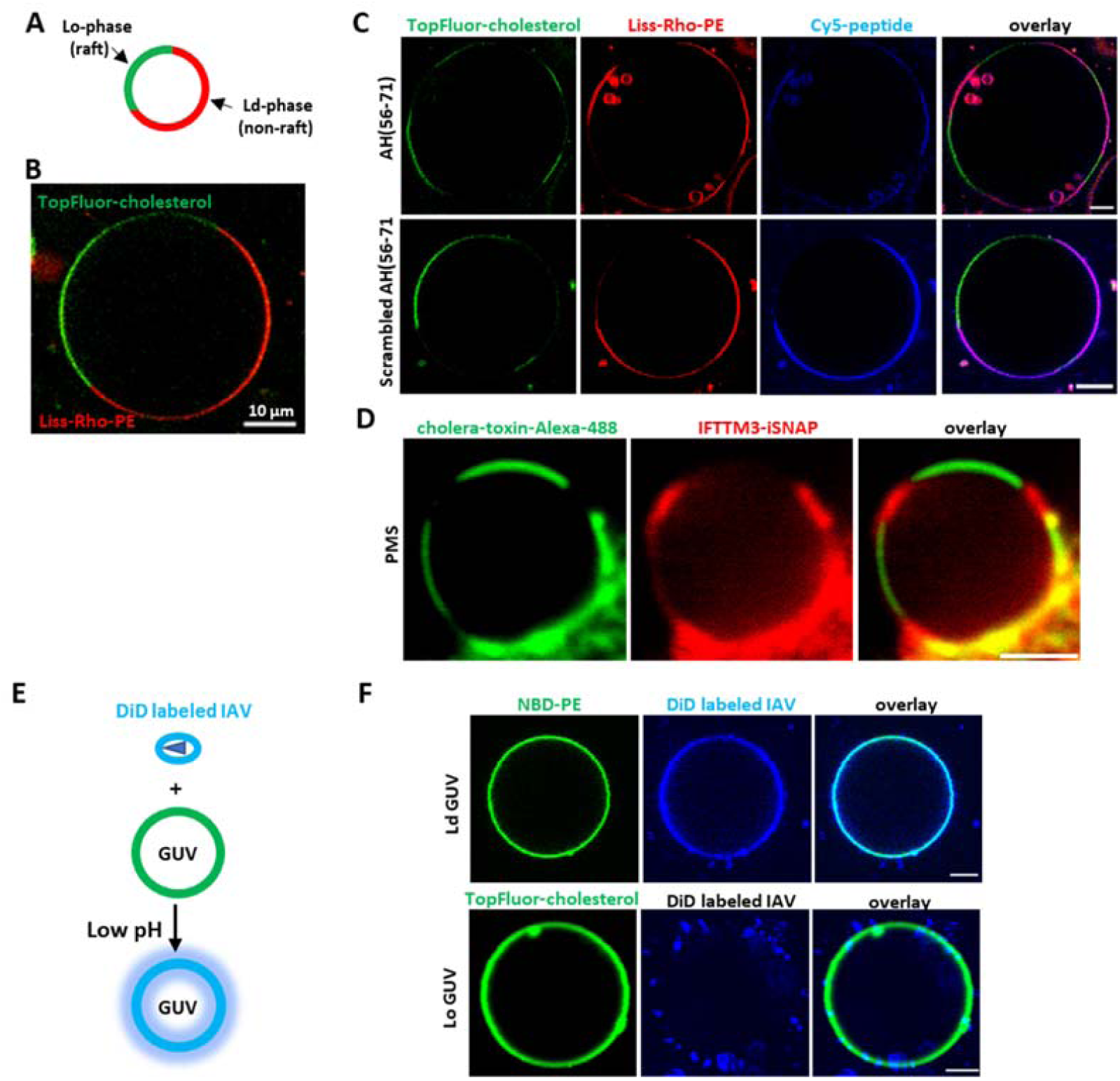
IFITM3 partitions into liquid-disordered membrane domains that support IAV fusion. (A) A diagram of phase-separated GUV. (B) A representative example of phase-separated GUV containing 33.3 mol % DOPC, 33.3 mol % SM, 32.4 mol % cholesterol, 0.5 mol % TopFluor-cholesterol (marker of Lo domain) and 0.5% Liss-Rho-PE (marker of Ld domain). (C) Phase-separated GUVs (33.3 mol % DOPC, 33.3 mol % SM, 32.4 mol % cholesterol, 0.5 mol % TopFluor-cholesterol and 0.5% Liss-Rho-PE) were incubated with 10 μM of Cy-5-labeled AH (56-71) or Scrambled AH(56-71) for 30 min and imaged. Scale bars 10 μm. (D) Plasma membrane spheres were prepared from IFITM3-iSNAP expressing A549 cells by cell swelling. GM1 was crosslinked with Cholera toxin B-AF488 (green) to mark the Lo phase and SNAP tag was stained with SNAP-cell 647-SIR (red). Scale bar 2 μm. (E) A diagram depicting lipid mixing between DiD-labeled IAV and GUV triggered by low pH leading to DiD dequenching. (F) Ld GUVs (top, 97.5 mol % DOPC, 2% GM1, 0.5% NBD-PE) or Lo GUVs (bottom, 66.6 mol % SM, 30.9 mol % cholesterol, 2% GM1, 0.5 mol % TopFluor-cholesterol) were mixed with DiD labeled IAV, lipids mixing was triggered by addition of citrate buffer to achieve the final pH of 5.0, and samples immediately imaged. Scale bars 5 μm. See also Figure S4.

We then asked whether the full-length IFITM3 protein similarly localizes to the Ld phase in cellular membranes. IFITM3 transits through the plasma membrane *en route* to endolysosomal compartments and is thus present on the cell surface (Jia et al., 2014). We thus used plasma membrane-derived spheres (Lingwood, Ries, Schwille, & Simons, 2008) to visualize IFITM3 partitioning into membrane rafts. We expressed IFITM3 with an internal SNAP-tag inserted at the same site as EGFP in IFITM3-iGFP (Suddala et al., 2019). SNAP-tagged IFITM3 was expressed in A549 cells and specifically labeled with a fluorescent substrate SNAP-Cell 647-SiR. Next, membrane spheres derived from the plasma membrane were obtained by cell swelling and phase separation within these spheres was induced by crosslinking a lipid raft marker GM1 with a labeled cholera toxin B subunit (Lingwood et al., 2008) (Figure 4D). The IFITM3-iSNAP partitioned to the Ld phase of membrane spheres, suggesting that the full-length IFITM3 is also localized to non-raft domains in living cells (Figure 4D).

Next, we sought to test whether the IFITM3 localization to Ld phase may be augment for its antiviral activity. To answer this question, we employed a lipid mixing assay between IAV labeled with a self-quenching concentration of the lipophilic dye DiD and GUVs. Here, addition of labeled IAV to GUVs followed by acidification should trigger lipid mixing which can be detected by the appearance of dequenched DiD in the GUV membrane (Figure 4E). Using phase-separated GUVs, we found that, upon triggering IAV-GUV fusion by exposure to low pH, DiD accumulated within the Ld domain <u>(Figure S4B)</u>, <u>although in the presence of IAV receptor ganglioside GM1, which dominantly accumulates in Lo domain (Figure S4C)</u>, suggesting that IAV lipid mixing occurred in a non-raft phase (Figure S4C). To test the possibility of DiD redistribution between Lo and Ld phases after viral fusion, single-phase GUVs were prepared. Low pH-induced IAV lipid mixing occurred very efficiently with Ld-phase GUVs (Figure 4F), whereas no lipid mixing was detected between IAV and Lo-phase GUVs, in spite of the efficient binding of labeled viruses to these vesicles (Figure 4F). Taken together, these results imply that IFITM3 concentrates in non-raft domains which appear to be the sites supporting IAV fusion in synthetic membranes.

### IFITM3 amphipathic helix increases lipid order and stiffens membranes

Since IFITM3 tends to localize to non-raft microdomains of membranes and induces negative curvature through its AH, we asked whether this helical domain affects other membrane properties, such as lipid order and membrane stiffness. Lipid order, which reflects the mobility of hydrocarbon tails of lipids (Vanblitterswijk, Vanhoeven, & Vandermeer, 1981), has been assessed using the lipophilic probe Laurdan. Laurdan is an environment-sensitive dye that intercalates between the hydrocarbon tails of lipids and exhibits a red-shift in the emission spectrum upon exposure to polar solvent, i.e., in lipid disordered domains. The Laurdan emission shift is parameterized by generalized polarization (GP), which is a normalized difference between Laurdan’ s emission at two wavelengths (Parasassi, De Stasio, Dubaldo, & Gratton, 1990) (Figure 5A). We thus used Laurdan to probe the effect of IFITM3 AH peptides on LUVs. Addition of either AH (56-71) or AH (59-68) peptide caused a strong positive shift in the Laurdan GP, indicating a marked increase in lipid order (Figure 5B). In contrast, the AH (59-68) F63Q mutant did not have a significant effect on GP, and the Scrambled AH (56-71) peptide only modestly altered GP. A detectable effect of the scrambled peptide on GP is likely due to its higher binding affinity to membranes (Figure 5B). These findings imply that IFITM3 AH specifically increases the lipid order.

**Figure 5.**
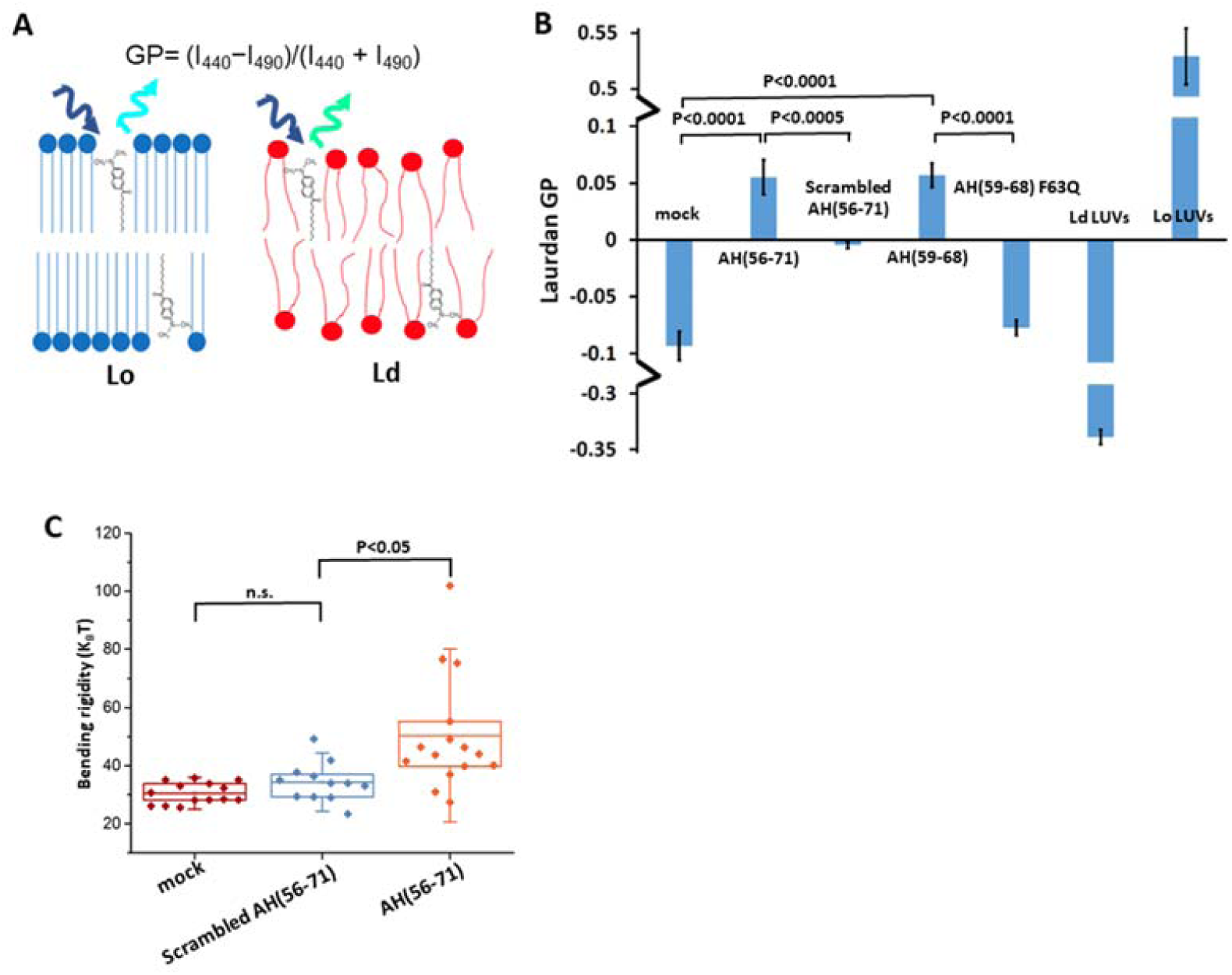
IFITM3 amphipathic helix increases lipid order and stiffens membranes. (A) A diagram illustrating the principle of Laurdan-based measurements of lipid order. (B)**M**embrane binding of IFITM3 AH induces lipid ordering. Two millimoles of LUVs (99.5 mol % POPC, 0.5 mol % cholesterol) were incubated with 25 μM Laurdan in the presence or absence of 40 μM of indicated IFITM3-derived peptide at 25°C. Two mM of Ld LUVs (99.5 mol % POPC, 0.5 mol % cholesterol, 37°C) and Lo LUVs (66.6 mol % SM, 33.4 mol % cholesterol, 25°C) were incubated with 25 μM Laurdan without addition of peptide. Laurdan fluorescence was then measured at 440 and 490 nm and the ratio was used to calculate the General Polarization (GP) of Laurdan using the equation GP= (I_440_−I_490_)/(I_440_ + I_490_). Data represent mean ± SD of three independent experiments. The statistical analysis used is the Student’ s t-test. (C) The membrane bending rigidity increases in the presence of IFITM3 AH peptide. Bending rigidity values were measured for POPC GUVs in 100 mM sucrose, 1mM HEPES at pH 7.4, incubated with DMSO (at final concentration of ∼0.1 v/v %; mock) or with AH(56-71) or Scrambled AH(56-71) in DMSO at final peptide concentration of 3 nM. Every data point corresponds to an individual vesicle. Boxes show the lower 25% and 75% quartile around the population mean value (middle line) and error bars indicate standard deviation.

It has been reported that lipid order correlates with the bending rigidity of cellular membranes (Steinkuhler, Sezgin, Urbancic, Eggeling, & Dimova, 2019). We therefore asked whether IFITM3 AH can affect the membrane’ s bending rigidity. The membrane bending rigidity was measured by fluctuation analysis of the thermally induced motion of GUV membrane, as described previously (Gracia, Bezlyepkina, Knorr, Lipowsky, & Dimova, 2010). Addition of AH (56-71) resulted in significant increase in the bending rigidity of the GUV membrane (Figure 5C). In contrast, the Scrambled AH (56-71) peptide did not have a significant effect on the bending rigidity. These important findings show that IFITM3 AH specifically stiffens lipid membranes and increases the lipid order.

### IFITM3 amphipathic helix alone is sufficient to inhibit membrane fusion

The ability of IFITM3 AH to induce negative curvature, increase the lipid order and stiffen the membrane implies that region is responsible for the protein’ s inhibitory effect on viral fusion. We hypothesized that AH interaction with the cytoplasmic leaflet of endosomal membranes is necessary and sufficient for the antiviral activity IFITM3. However, testing this hypothesis *in vitro* using reconstituted IFITM3 peptides is not feasible. This is because incoming viruses do not come into contact with the N-terminal cytoplasmic region of IFITM3 that encompasses the AH, whereas reconstituted IFITM3 and IFITM3 AH localize to the external leaflet of endosomes (Figure 1A) that comes in contact with exogenously added viruses. To fulfill the topological requirement for virus/IFITM3 positioning across a target membrane, using exogenously added peptides, we resorted to exocytosis (Figure 6A). Secretory vesicle fusion with the plasma membrane affords an easy access to the external (trans) leaflet of a target membrane. We used glucose-stimulated insulin secretion as a model to study the effect of IFITM3 on SNARE-mediated vesicle fusion (Figure 6A). INS-1E cells secrete insulin upon stimulation with high glucose (Merglen et al., 2004). To increase the sensitivity of detection of secreted insulin, INS-1E cells were transduced with a vector expressing a proinsulin-luciferase fusion protein (Burns et al., 2015), which can be detected by luciferase activity following exposure to a high glucose solution (Figure 6B). When cells were pretreated with AH (59-68), which effectively bound to the plasma membrane (Figure S5A), high glucose stimulation resulted in significantly reduced insulin secretion. In contrast, the AH (59-68) F63Q peptide had no effect on insulin secretion, even when added in a higher concentration than the wild-type peptide (Figure 6B, Figure S5B). To test whether inhibition of secretion was related to the IFITM3 AH-mediated negative membrane curvature, we pretreated cells with LPC, which induces positive curvature. LPC promoted insulin secretion, in agreement with the previous study (Amatore et al., 2006) (Figure 6C). Importantly, LPC added with IFITM3 AH counteracted the suppression of insulin secretion by this peptide. Taken together, these results imply that IFITM3 AH is sufficient to inhibit membrane fusion and that this inhibition is dependent on its ability to induce negative membrane curvature.

**Figure 6.**
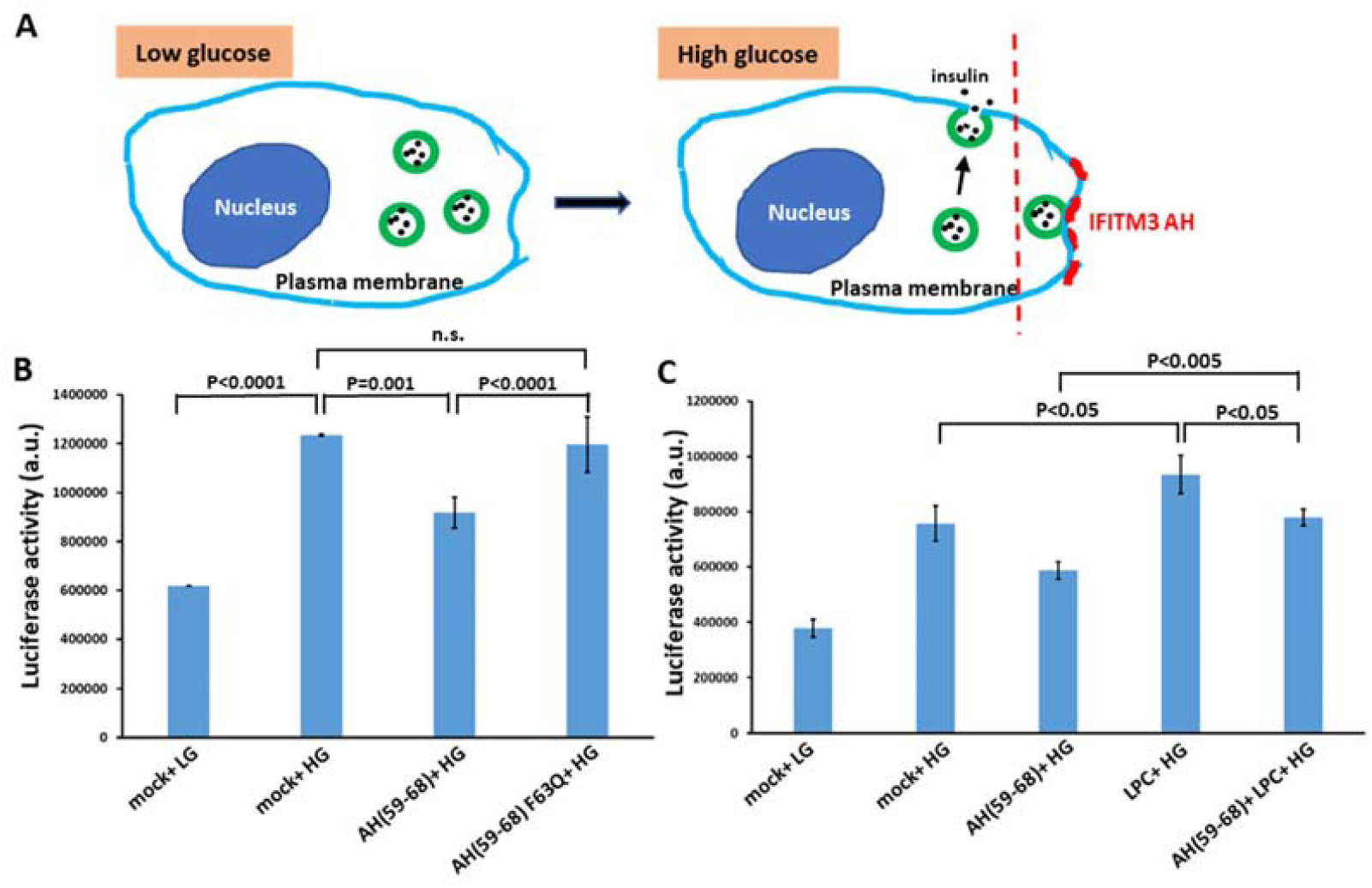
IFITM3 amphipathic helix inhibits membrane fusion. (A) A diagram illustrating glucose-stimulated insulin secretion and its block by IFITM3 AH. (B) IFITM3 AH peptide inhibited glucose-stimulated insulin secretion of INS-1E cells. INS-1E cells transduced with proinsulin-luciferase fusion construct were preincubated for 5 min with 5 μM AH(59-68) or 10 μM AH(59-68) F63Q or same volume of DMSO (mock) and then stimulated for 20 min with high glucose (HG, 20 mM) buffer or with low glucose (LG, 2.8 mM) as control. Luciferase activity was determined by adding the coelenterazine substrate to the supernatant and reading on a Luminescence counter. Data represent mean ± SD of the results of three independent experiments. The statistical analysis used is the Student’s t-test. (C) As in (B), but for samples preincubated for 5 min with AH peptide or DMSO supplemented with 20 µM LPC. See also Figure S5.

To further investigate the link between membrane curvature and antiviral activity of IFITM3, we asked whether the IFITM3 AH can be replaced with unrelated amphipathic helixes of other proteins, such as M2 of IAV (M2 AH). The M2 AH has been reported to induce negative curvature and increase lipid order during IAV budding from the plasma membrane (Martyna et al., 2017; Rossman, Jing, Leser, & Lamb, 2010). We thus replaced the IFITM3 AH with M2 AH in the context of an mTFP1 tagged protein (abbreviated IFITM3-M2 AH-imTFP1, Figure 7A). We also constructed a scrambled IFITM3 AH mutant (IFITM3-Scrambled AH-imTFP1) as a control (Figure 7A). When overexpressed in HEK 293/17 cells, wild-type IFITM3 significantly inhibited the infection of the mCherry-expressing IAV (Figure 7B, C). As expected, the scrambled IFITM3 AH mutant exhibited a markedly attenuated antiviral activity against IAV. Importantly, IFITM3 chimera with the M2 AH showed strong antiviral activity, without significantly affecting cell viability (Figure S5C). The demonstration that IFITM3 AH can be replaced by other negative curvature-inducing and lipid-ordering AHs further confirms that IFITM3 inhibits viral fusion by modulating the proprieties of endosomal membrane through its amphipathic helix.

**Figure 7.**
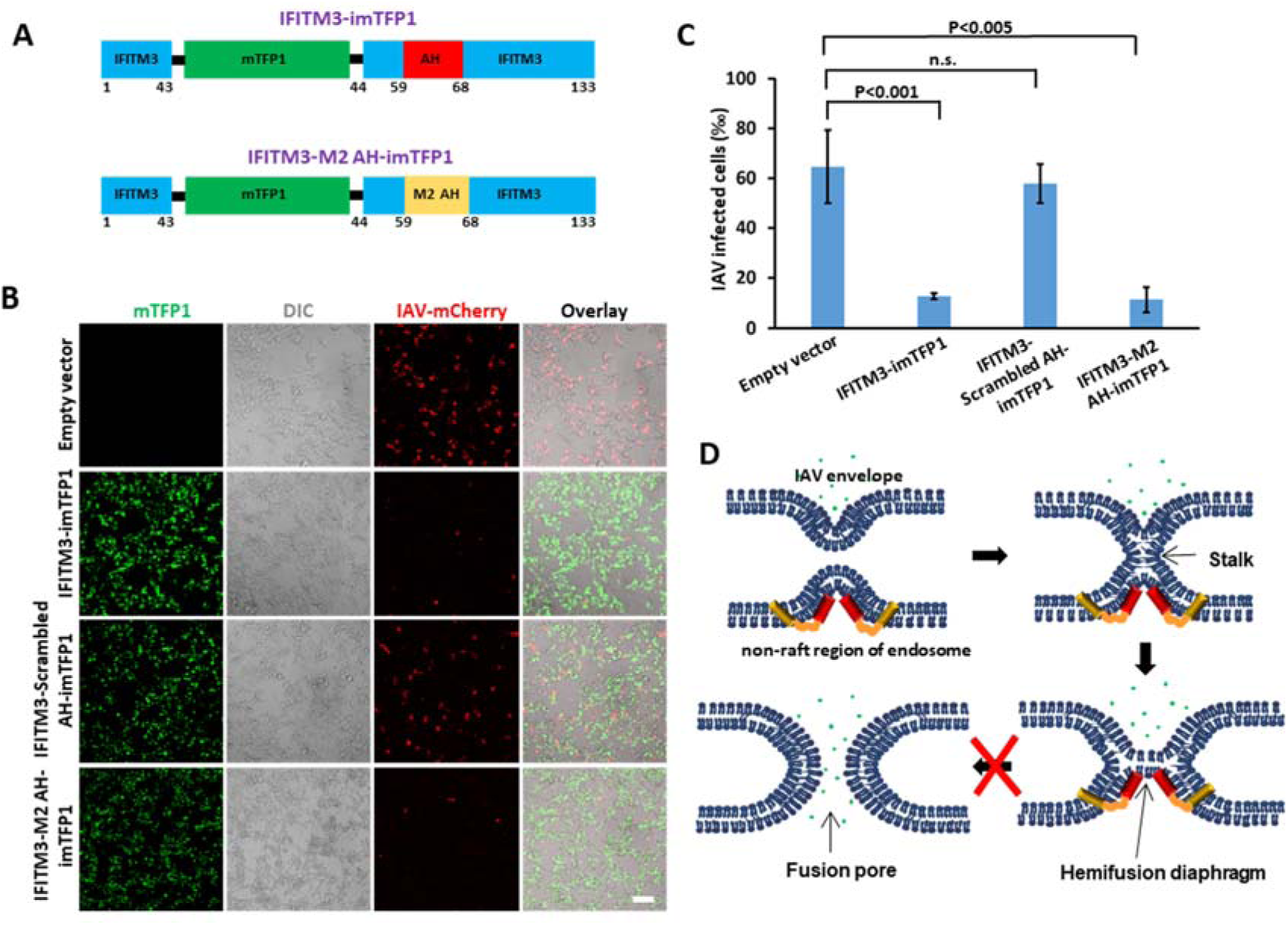
IFITM3 amphipathic helix can be replaced by other negative curvature-inducing and lipid-ordering amphipathic helices. (A) Illustration of the internal mTFP1-tagged IFITM3 AH chimera with the Influenza virus M2 protein AH. (B) Infection assay of HEK 293T/17 cells expressing wild-type IFITM3-imTFP1, its scrambled mutant or the M2 AH chimera using mCherry-expressing IAV. Sale bar 100 μ m. (C) Quantification of IAV infection shown in (B). Data represent mean ± SD of the results of three independent experiments. The statistical analysis used is the Student’s t-test. Total number of cells counted are: Empty vector (3813), IFITM3-imTFP1 (3877), IFITM3-scrambled AH-imTFP1 (3525), IFITM3-M2 AH-imTFP1 (4244). (D) Working model of IFITM3-mediated inhibition of IAV entry. IFITM3 localizes to liquid disordered subdomains of endosomal membrane, which are the sites supporting IAV fusion, and stabilizes the hemifusion diaphragm by inducing negative membrane curvature and increasing lipid ordering and membrane stiffness. This prevents the formation of a fusion pore in the hemifusion intermediate, thus trapping IAV at a “dead-end” hemifusion stage. See also Figure S5.

## Discussion

A remarkably broad-spectrum of enveloped viruses that are restricted by IFITM proteins suggests a universal mechanism for their antiviral activity. This mechanism may involve modifying the properties of the host cell membranes to disfavor fusion pore formation and thus trap viral fusion at a “dead-end” hemifusion stage (Desai et al., 2014; Suddala et al., 2019). Here, we provided direct evidence for this mechanism by showing that lipid bilayer-reconstituted recombinant IFITM3 induces unfavorable negative curvature and increases the lipid order and membrane stiffness. Importantly, we found that these effects on lipid bilayers *in vitro* are linked to the IFITM3’ s ability to inhibit membrane fusion in cells. Specifically, we showed that the incorporation of IFITM3 AH into the external leaflet of the plasma membrane inhibits exocytosis (Figure 6B). Although exocytic fusion is the topological opposite of viral fusion, it provides a convenient model system for assessing the effects of exogenously added amphipathic peptides and lipids (Amatore et al., 2006; Burns et al., 2015).

According to the stalk-pore hypothesis (Chernomordik, Melikyan, & Chizmadzhev, 1987; Kozlov, Leikin, Chernomordik, Markin, & Chizmadzhev, 1989), membrane fusion proceeds through a series of highly curved intermediates – stalk, hemifusion and fusion pore – that are characterized by different net curvatures and thus respond differently to changes in lipid shape/composition. The formation of a lipid stalk involves a local disruption and bending of contacting membrane leaflets into a net negative curvature structure (Figure 7D). The hemifusion intermediate is also a net negative curvature structure, whereas the fusion pore possesses a net positive curvature (Figure 7D) (Chernomordik & Kozlov, 2003). Thus, the presence of lipids that induce positive curvature, such as LPC, in the contacting leaflets blocks the stalk formation, whereas their incorporation into the distal leaflet of a target membrane, promotes rupture of the hemifusion diaphragm and fusion pore formation (Amatore et al., 2006; Chernomordik, Frolov, Leikina, Bronk, & Zimmerberg, 1998; Chernomordik et al., 1987; Stiasny & Heinz, 2004). In contrast, fusion pore formation is impaired by lipids inducing negative curvature, such as oleic acid (OA) (Chernomordik, Leikina, Frolov, Bronk, & Zimmerberg, 1997). Thus, spontaneous curvature of lipids is an essential determinant of membrane hemifusion and fusion mediated by cellular and viral proteins (Chernomordik & Kozlov, 2003).

Our experiments demonstrate that, similar to OA, IFITM3 AH induces negative curvature of GUVs manifested in inward budding and ILV formation. Thus, in the context of IFITM3 expressed in cells, AH inserted into the cytoplasmic leaflet of an endosomal membrane is expected to stabilize the hemifusion diaphragm formed by incoming viruses and thus prevent the fusion pore formation (Figure 7D). In addition, the increased membrane order and stiffness induced by IFITM3 are expected to further disfavor membrane fusion (Chesarino et al., 2017; John et al., 2013; Li et al., 2013; Lin et al., 2013).

Our finding that IFITM3 induces negative curvature and inward budding in GUVs is consistent with the observation that its overexpression in cells robustly induced the formation of large multivesicular bodies that are full of IFITM3-containing intralumenal vesicles (Amini-Bavil-Olyaee et al., 2013). These results also support the “fusion decoy” model, according to which excess of intralumenal vesicles in IFITM3-expressing cells favors non-productive viral fusion with intralumenal vesicles, instead of fusion with the limiting membrane of endosomes (Desai et al., 2014). The above considerations suggest that a potent inhibition of diverse enveloped viruses by IFITM3 may involve a combination of modulation of membrane properties and generation of an excess of decoy vesicles in endosomes.

Lipid raft domains have been proposed to be the entry sites of viruses due to the raft localization of certain signaling proteins and receptors (Chazal & Gerlier, 2003). However, direct evidence supporting this model is lacking due, in part, to the dynamic nature of raft micro- and nano-domains in living cells (Rajendran & Simons, 2005b). In addition, the rigid nature of lipid raft domains is expected to disfavor membrane fusion. Interestingly, it has been reported that the boundaries between ordered and disordered lipid domains, which are characterized by high line tension (energy per unit length), are the predominant sites of HIV-1 fusion (Yang et al., 2017). However, such apparent preference for domain boundaries is not universal, since IAV lipid mixing occurs efficiently with Ld-phased GUVs lacking lipid rafts or phase boundaries (Figure 4F). No lipid mixing could be detected with Lo GUVs. A correlation between the apparent IAV fusion site and the IFITM3 localization to Ld phase may offer a clue regarding the potential mechanism of virus resistance. It is tempting to speculate that IFITM3-resistant viruses, like Lassa virus, may localize to and fuse with lipid raft domains devoid of this restriction factor.

It is worth pointing out that AH is present in IFITM1 and IFITM2 proteins and is highly conserved across vertebrates (Chesarino et al., 2017; Z. Zhang et al., 2012). It is therefore very likely that the AH region is key to the antiviral activity of all three human restriction factors. Clearly, this novel proximity-based antiviral mechanism is dependent on proper trafficking of IFITMs to ensure their concentration at the sites of virus entry (Li et al., 2013; Suddala et al., 2019). Accordingly, incorporation of IFTIMs into virions ensures their presence at the site of fusion and effectively inhibits infection of nearly all viruses, including those that are otherwise resistant to IFITM restriction when expressed in target cells (Appourchaux et al., 2019; Suddala et al., 2019; Tartour et al., 2017)

Our findings reveal a universal defense mechanism employed by cells to effectively ward off invading enveloped viruses through modification of the cytoplasmic leaflet of cellular membranes. The IFITM AH-mediated increases in negative curvature and stiffness of the cytoplasmic leaflet disfavor the transition from hemifusion to full fusion and thereby block entry of diverse enveloped viruses. This restriction mechanism informs us of novel antiviral strategies targeting the cytoplasmic leaflet of cell membranes.

## Materials and Methods

### Cell lines, plasmids and reagents

HEK293T/17 and A549 cells were obtained from ATCC (Manassas, VA) and were maintained in Dulbecco’s Modified Eagle Medium (DMEM, Corning, Corning, NY) containing 10% heat-inactivated Fetal Bovine Serum (FBS, Atlanta Biologicals, Flowery Branch, GA) and 1% penicillin/streptomycin (Gemini Bio-products, West Sacramento, CA). For HEK 293T/17 cells, the growth medium was supplemented with 0.5 mg/ml G418 (Genesee Scientific, San Diego, CA). INS-1E cells were obtained from Addexbio Technologies (San Diego, CA, USA) and were maintained in RPMI-1640 Medium (Addexbio), supplemented with 10% heat-inactivated FBS, 10 mM HEPES, and 55 μ M β -mercaptoethanol (Invitrogen, Carlsbad, CA). pET28a vector was a gift from Dr. Baek Kim (Emory University). pQCXIP was purchased from Clontech (Mountain View, CA). Proinsulin-NanoLuc in pLX304 vector (plasmid # 62057), psPAX2 packaging plasmid (plasmid #12260) and pMD2.G envelope plasmid (plasmid #12259) were from Addgene (Watertown, MA). N-terminally Cy5-labeled peptides were synthesized and purified to > 95% purity by Gen-Script (Piscataway, NJ). Peptide sequences used were: wt-long, Cy5-DHVVWSLFNTLFMNPC; scrambled peptide, Cy5-LVWHMFLSDNFTNPV; wt-short, Cy5-CVWSLFNTLFM and F63Q short, Cy5-CVWSLQNTLFM. The fluorescently labeled, palmitoylated IFTIM3 peptide (FITC-Ahx-DHVVWSLFN TLFMNPC(PAL)CLGF) was chemically synthesized on Rink Amide MBHA resin using Fmoc solid phase peptide synthesis, and Fmco-Cys(Mmt)-OH and Fmoc-Ahx-OH, among other Fmoc-protected amino acids. After chain assembly, the Fmoc group of the N-terminus was removed and extended by the linker amino acid Ahx, followed by the coupling of fluorescein isothiocyanate (FITC, 2 eq.) and DIPEA (2 eq.) in DMF (6 mL) at room temperature overnight. The monomethoxytrityl (Mmt) group was removed with diluted TFA (2%, DCM) and scavengers (10% TIPS), followed by the addition of palmitic anhydride (10 eq.) dissolved in DCM and DIEA (20 eq.) for 3 h. After cleavage from the resin and precipitation with cold diethyl ether, the crude peptide was purified by revered phase HPLC and its molecular mass verified by ESI-MS.

mCherry-expressing IAV PR/8/34 was a gift from Dr. Luis Martinez-Sobrido (University of Rochester). The following lipids were purchased from Avanti Polar Lipids (Alabaster, AL): 1-oleyl-2-palmitoyl-sn-glycero-3 phosphocholine (POPC), 1,2-dioleoyl-sn-glycero-3-phosphocholine (DOPC), 1-stearoyl-2-hydroxy-sn-glycero-3-phosphocholine (18:0 Lyso PC), N-stearoyl-D-erythro-sphingosylphosphorylcholine (18:0 SM), 23-(dipyrrometheneboron difluoride)-24-norcholesterol (TopFluor® Cholesterol), cholesterol (plant derived), and 1,2-dioleoyl-sn-glycero-3-phosphoethanolamine-N-(lissamine rhodamine B sulfonyl) (18:1 Liss-Rho-PE).

### Protein expression and purification

IFITM3 gene was PCR-amplified and cloned into the pET28a vector to produce recombinant proteins fused to Strep-tag II (WSHPQFEK). Point mutations and deletions were introduced by site-directed mutagenesis. The IFITM3 constructs were transformed into Escherichia coli Rosetta(tm) 2 (DE3) pLysS Singles(tm) Competent Cells (MilliporeSigma, Burlington, MA) to overexpress the proteins. The bacteria were cultured in Terrific broth medium at 37°C. Protein expression was induced by the addition of 0.3 mM IPTG when OD600 was 1.0 and culture for additional 20 h at 16°C. Cells were harvested after centrifugation at 2,500 ×g for 30 min, resuspended and sonicated in HND buffer (25 mM HEPES, pH 7.4, 150 mM NaCl, 1 mM DTT) supplemented with cOmplete EDTA-free Protease Inhibitor Cocktail (Roche, Basel, Switzerland). The lysate was then centrifuged at 4° C in the SW32Ti rotor with a speed of 30,000 rpm for 1 h to pellet the membrane fraction using an Optima L-90K Ultracentrifuge (Beckman Instruments, Brea, CA). The pellet was resuspended in HND buffer and bulk membrane proteins were extracted by adding 2% ANAPOE-X-100 (Anatrace, Maumee, OH) and incubating for 1 h at 4° C, followed by a 1 h-centrifugation at 30,000 rpm. The supernatant was loaded on Strep-Tactin resin (QUIGEN, Hilden, Germany), incubated for 3 h at 4°C, and the column was washed twice with 15 ml HND buffer supplemented with 0.1% ANAPOE-X-100. Proteins were eluted by adding 10 ml HND buffer supplemented with 0.1% ANAPOE-X-100 and 2.5 mM desthiobiotin and then concentrated using an Amicon Ultra 5,000 MW cutoff filter (Millipore, Billerica, MA). Protein purity was assessed by SDS-PAGE and Coomassie Blue staining. For purification of wild-type and mutant transmembrane-truncated IFITM3, cell lysate was cleared by centrifugation at 4°C with a speed of 18,000 rpm for 45 min after sonication, and the supernatant was directly loaded on Strep-Tactin resin and incubated for 3 h at 4°,washed and eluted with the buffer used for purification of the full-length IFITM3 except that ANAPOE-X-100 was omitted.

### Large Unilamellar Vesicles

Lipids (99.0 mol % POPC, 0.5 mol % cholesterol, 0.5 mol % Liss-Rho-PE) were mixed in a glass tube and dried down to a film under a gentle stream of Argon, followed by further drying under a vacuum for 30 min. Next, the lipid film was hydrated and resuspended with HND buffer to a final lipid concentration of 10 mM. Large unilamellar vesicles (LUVs) were formed from the lipid suspension by ten freeze-thaw cycles using liquid nitrogen and room temperature water bath. Uniform-sized LUVs were formed by extruding through polycarbonate filters with 100-nm pore size (Avanti Polar Lipids, Alabaster, AL) 11 times.

To reconstitute IFITM3, preformed liposomes and purified IFITM3 (molar protein to lipid ratio, 1:500) were mixed with 0.1% Triton X-100 at an effective detergent to lipid ratio of ∼1 and incubated for 1h at 4 °C. Triton X-100 was then removed by adding BioBeads SM-2 absorbent beads (BioRad) at a Bio-Beads/Triton X-100 ratio of 10 (wt/wt) in five portions during the course of hour, and incubating overnight after the final addition of beads. Insoluble protein aggregates were pelleted by centrifugation of samples in an Eppendorf microcentrifuge (10 min, 16,000× g).

### Liposome co-floatation assay

Peptide binding to LUVs was measured by mixing 204 μ l proteoliposomes with 600 μ l of a 67% sucrose solution, bringing the final sucrose concentration to 50%. The mixture was transferred to a clear polycarbonate ultracentrifuge tube and overlaid with two layers consisting of 10.2 ml of 25% sucrose and 1 ml of 5% sucrose. After centrifugation at 30,000 rpm (4 ° C for 3 h) in the SW45Ti rotor (Beckman Instruments), eight 1.5-ml fractions were collected from the top of a gradient. 30 μ l aliquots from each fraction were analyzed by Western blotting with IFITM3 antibody (N-term, Abgent, San Diego, CA).

### Trypsin cleavage assay

To determine the accessibility of LUV-reconstituted peptides, 20 μ l proteoliposomes was incubated with 0.6 μ g trypsin (TPCK-Treated, Sigma-Aldrich) at 37° C for 30 min. Next, 95° C pre-heated SDS sample buffer (50 mM Tris-HCl, pH 6.8, 2% SDS, 10% glycerol, 1% β -mercaptoethanol, 12.5 mM EDTA, 0.02 % bromophenol blue) was added to stop the reaction. The samples were analyzed by SDS-PAGE and stained with Coomassie blue.

### Liposome co-sedimentation assay

Peptides were sonicated in water bath for 30 min and possible aggregates were removed by spinning down at 20,000 × g for 5 min. Two mM of LUVs made of 99.0 mol % POPC, 0.5 mol % cholesterol, 0.5 mol % Liss-Rho-PE in PBS were incubated at room temperature with 20 μM IFITM3(1-108)-iEGFP protein or 40 μM of Cy5-labeled peptides for 10 min in a 100 μl reaction volume. The mixture was then diluted with PBS and centrifuged at 30,000 rpm for 30 min at 20 °C. Fluorescence of supernatant was measured on a SpectraMax i3x microplate reader (Molecular Devices, CA, USA). The binding efficiency of proteins/peptides was decided by reduction of EGFP or Cy5 fluorescence in supernatant and normalized by comparing with the fluorescence of input samples.

### Giant Unilamellar Vesicles

GUVs were prepared from a 5 mM solution of lipids in chloroform (99 mol % POPC, 0.5 mol % cholesterol and 0.5 mol % of Liss-Rho-PE). Fifty μl of the lipid solution was spread onto a 10 cm^2^ area of the conductive side of each of two indium-tin oxide (ITO) coated slides (70-100 Ω, Delta Technologies, Stillwater, MN), allowed to evaporate and kept under vacuum for 1 hr. Electroformation chambers were constructed by sandwiching a 1 mm spacer between two lipid-coated slides. Next, the chamber was filled with 0.1 M sucrose, 1 mM HEPES, pH 7.0, and a 10 Hz sine wave voltage (1 V peak-to-peak) was applied across the chamber for 3 h using a function generator. GUVs were gently collected with a pipette and used immediately.

To prepare phase-separated GUVs, we used a mixture of 33 mol % DOPC, 33 mol % SM, 33 mol % cholesterol, 0.5% TopFluor-cholesterol, 0.5% Liss-Rho-PE. Lipids were dissolved in 50 µ l chloroform/methanol (9:1) in a glass tube and placed in a preheated block at 60 ° C for 1 min after gentle vortexing and centrifugation. The mixture was then immediately deposited and spread over 2 preheated ITO coverslips, and the solvent was evaporated at 60 ° C. After ITO coverslips dried out, they were immediately placed into a vacuum chamber for 1h to remove residual solvent. All these steps were done in a timely manner to minimize possible lipid oxidation. GUV electroformation was carried out in 0.1 M sucrose, 2 mM DTT, 1 mM HEPES, pH 7.0 at 60 ° C for 4 h. Collected GUVs were cooled to room temperature and used immediately. For imaging of GUVs with IFITM3 peptides, peptides were prepared as 2 mg/ml stock solutions in DMSO. Twenty μ l GUVs were diluted in 30 μ l of a hypertonic buffer containing160 mM sucrose, 1 mM HEPES, pH 7.0 to slightly deflate GUVs. Next, 2 μ l of peptide diluted with 98 μ l Hank’ s Balanced Salt Solution (Corning, NY, USA) was added to and mixed with GUVs, yielding the final concentration of peptide of 10 μ M. The mixture of GUVs and peptide was immediately added to an 8-well chambered coverslip pretreated with a BSA solution (2 mg/ml) for 30 min to attach GUVs to the bottom of the chamber. GUVs were allowed to sediment for 10 min before imaging on a Zeiss LSM880 laser scanning confocal microscopes, using a 63x/1.4NA oil-immersion objective. Purified wild-type and mutant transmembrane-truncated IFITM3 proteins were imaged the same way except that the final concentration of proteins was 20 μM. Quantification of inward budding of GUVs was performed by counting all GUVs with diameter above 2 μm, regardless of whether they encompassed intraluminal vesicles containing aqueous dye (fluorescein or EGFP). Images of at least 2 randomly selected fields of view were acquired using Z-stacks separated by 1 µ m. For each condition, at least two independent experiments were performed. Statistical analysis was done using the Student’ s t-test.

GUVs containing IFITM3 protein were prepared through dehydration of IFITM3-containing LUVs, as previously described (Girard et al., 2004). Briefly, IFITM3-containing LUVs were prepared, as above, added dropwise onto indium-tin oxide coated slides, and dehydrated under vacuum overnight. The lipids were rehydrated and electroformed, as described above at room temperature for 6 h.

### Plasma membrane spheres

Plasma membrane spheres were prepared as previously described (Lingwood et al., 2008). Briefly, A549 cells transduced with IFITM3-iSNAP/pQCXIP expression vector were seeded on 8-well chambered coverslip. 24 h later cells were washed and incubated for 18 hr in PMS buffer (1.5 mM CaCl_2_, 1.5 mM MgCl_2_, 5 mM HEPES pH 7.4, 1 mg/ml glucose in 1x PBS) at 37 ° C. The lipid ordered phase was visualized through GM1 crosslinking by incubating the PMS in 10 µg/ml of Alexa-488 labeled cholera-toxin subunit B (Invitrogen) for 2 h at 37°C and IFITM3-iSNAP was stained with 3 µM SNAP-cell 647-SIR (New England Biolabs) at the same time. PMS were then washed and imaged at room temperature on a Zeiss LSM880 laser scanning confocal microscopes, using a 63x/1.4NA oil-immersion objective.

### Lipid mixing between influenza virus and GUVs

Virus labeling was performed essentially as described previously (Suddala et al., 2019). Briefly, a 100 µ l aliquot of the 2 mg/ml Influenza A/PR/8/34 virus stock (Charles River, CT, USA) was thawed at room temperature and diluted using 50 µ l isotonic 145 mM NaCl/50 mM HEPES (pH 7.4) buffer. To label the virus membrane, 1.5 µ l of DiD dye (Invitrogen, 10 mM stock solution in DMSO) was quickly injected into a mixture during mild vortexing to final concentration of 100 µ M. The tube was closed, wrapped with aluminum foil and agitated at the lowest speed setting of a vortex for 1 hour at room temperature. The labeled viruses were purified from excess dyes on a Nap-5 gel filtration column (GE Healthcare) that was equilibrated with 50 mM HEPES, pH 7.4, 145 mM NaCl at room temperature. The fractions containing labeled viruses were passed through a 0.45 μ m filter to remove any large lipid and/or virus aggregates. To test the labeling efficiency, 10 µ l of the purified viruses were diluted to 1000 µl in PBS, lysed with 0.5% TX-100 (final concentration) and the extent of DiD dequenching was measured in a plate reader. A more than 20-fold increase in the DiD signal upon lysis has been found to give good results in lipid mixing experiments. The labeled virus was aliquoted into tubes, flash-frozen, and stored at -80 ° C until use. The virus lipid mixing assay was performed by mixing 20 µ l of the labeled virus with 30 µl GUVs, followed by addition of 150 µl of 100 mM citrate buffer (pH 4.8) supplemented with 100 mM MES to achieve the final pH of 5.0. The resulting mixture was transferred into a BSA-pretreated 8-well coverslip and imaged on a Zeiss LSM880 laser scanning confocal microscope, using a 63x/1.4NA oil-immersion objective.

### Lipid order measurement

To assess the lipid order, 2 mM of LUVs were mixed with 25 μM of Laurdan dye (Invitrogen) and 40 μM of indicated IFITM3-derived peptide in a total volume of 50 μl. Fluorescence was measured on a SpectraMax i3x microplate reader using a 355 nm excitation filter and recording fluorescence emissions at two wavelengths centered at 440 and 490 nm. Laurdan general polarization (GP) was calculated using the equation: GP = (I_440_− I_490_)/(I_440_ + I_490_), where I_440_ and I_490_ are fluorescence intensities at 440 nm and 490 nm, respectively.

### Membrane bending rigidity measurement

GUVs were grown on a polyvinyl alcohol (PVA) film as described previously (Weinberger et al., 2013). Briefly, 50 µl of 5% w/v (145000 g/mol) PVA (Merck) solution in water was spread on a cleaned (rinsed in ethanol and double distilled water) glass slide to form a thin film. The PVA film was dried kept in the oven at 50° C for 30 minutes. Five µ l of 2 mM lipid solution (80 mol % POPC, 20 mol % cholesterol) in chloroform was spread on the PVA film and the solvent was evaporated in vacuum for 1 hour. An observation chamber with a volume of about 700 µ l was assembled using a Teflon spacer sandwiched between two glasses with the lipid film facing inward, and the lipid films were hydrated in solution of 100 mM sucrose (Sigma), 1 mM HEPES (Sigma) at pH 7.4. After swelling for 30 minutes, GUVs were harvested and used right away. An aliquot of 0.1 μ l DMSO solution with or without the peptide was carefully pipetted into 100 μ l GUV suspension and slowly agitated to ensure mixing. Subsequently, 30 μl of the final solution was placed on a BSA coated glass coverslip. A few percent of the solution was left to evaporate for three to five minutes to ambient air and then closed in an observation chamber. Bending rigidity analysis was performed either on vesicles with added DMSO only (control) or DMSO with IFITM3 at final concentration of the peptide of 3 nM. Membrane bending rigidity was measured by fluctuation analysis of the thermally induced motion of the membrane, as described previously (Gracia et al., 2010). Experiments were performed on an Axio Observer D1 microscope (Zeiss, Germany) using a 40× objective in phase contrast mode. Imaging was performed using a low noise liquid-cooled digital camera pco.edge 5.5 a total of 1000-2000 snapshots per vesicle were acquired with exposure time of 200 µs at 10 frames per second.

### Virus infection assay

Luciferase-expressing pseudoviruses were produced by transfecting HEK293T/17 cells using JetPRIME transfection reagent (Polyplus-transfection, SA, NY), as described previously (Francis, Marin, Shi, Aiken, & Melikyan, 2016). Briefly, HEK293T/17 cells grown in a 100-mm dish were transfected with 2.5 μg of pCAGGS plasmid expressing H1N1 HA and NA proteins, 4 μg NL4-3R^-^E^-^Luc, and 1 μg pcRev. The transfection medium was replaced with fresh DMEM/10% FBS after 12 h, and cells were cultured for additional 36 h, after which time, the virus-containing culture medium was collected, passed through a 0.45-μm filter, and concentrated 10x using Lenti-X Concentrator (Clontech, Mountain View, CA). Following an overnight concentration with Lenti-X, virus was precipitated by centrifuging at 1439xg for 45 min at 4 °C, resuspended in DMEM without phenol red or FBS, and stored at -80 °C. Infection assays were performed using HEK 293T/17 cells transfected with indicated IFITM3 constructs in pcDNA3.1(+) vector using JetPRIME transfection reagent. At 24 h post-transfection, pseudoviruses (0.1 ng of p24) were spinoculated onto cells at 1,500×g, 4°C for 30 min, and cells were cultured for 24 h. Luciferase activity was determined by adding Bright-Glo Luciferase substrate (Promega, WI, USA) and reading with a TopCount NXT Luminescence counter (PerkinElmer, Waltham, MA,USA).

For infection of mCherry-expressing IAV PR8 virus, HEK 293T/17 cells were seeded on 8-well chambered coverslip 24 h before transfection with indicated IFITM3 constructs in pQCXIP. At 12 h post-transfection, mCherry-expressing IAV PR8 virus (2.5×10^4^ PFU/ml) was spinoculated onto cells at 1,500×g, 4°C for 30 min, and cells were cultured for 12 h and imaged on a Zeiss LSM880 laser scanning confocal microscope, using a 20x objective.

### Insulin secretion assay

An insulin secretion assay was performed, as described previously (Burns et al., 2015) with some modifications. Briefly, A lentivirus vector expressing luciferase fusion protein was produced using a second-generation viral packaging system. 4 μ g of pLX304 vector containing the fusion construct Proinsulin-NanoLuc, 2 μ g of psPAX2 packaging plasmid, 2 μg of pMD2.G envelope plasmid were used to transfect a 10 cm dish of HEK 293T/17cells using JetPRIME transfection reagent. Virus was harvested at 48 hours post-transfection and passed through 0.45 μ m cellulose acetate filters prior to use. Viruses were spun onto INS-1E cells at 800 xg for 1 hour at 30 ° C. After 24 hours at 37 ° C in 5% CO_2_ in the presence of virus, the medium was placed in fresh growth media with 5 μ g/mL blasticidin (Invitrogen) for 3 days to select for infected cells. For insulin secretion assays, INS-1E cells were plated in 96-well plates and cultured overnight at 37 °C in 5% CO_2_. Cells were washed once with PBS and incubated for 1 hour at 37 °C in sterile, 0.45 μm-filtered Krebs Ringer Buffer (KRB) containing 138 mM NaCl, 5.4 mM KCl, 2.6 mM MgCl_2_, 2.6 mM CaCl_2_, 5 mM NaHCO_3_, 10 mM HEPES and 5 g/L BSA (Sigma), supplemented with 2.8 mM glucose. The cells were then incubated with 2 µ M of indicated IFITM3 peptides in KRB for 5 min and then stimulated for 20 mins in 100 μ l of fresh KRB with 20 mM glucose. For the time course study, after incubating the cells for one hour in 2.8 mM glucose KRB, the buffer was changed every 5 minutes to fresh, prewarmed to 37 °C KRB containing either 2.8 mM glucose (for the first time point) or 20 mM glucose (for all subsequent time points). The luciferase signal in samples was determined by adding the coelenterazine substrate (NanoLight, Pinetop, AZ) to the supernatant to a final concentration of 10 µM and reading on a TopCount NXT Luminescence counter.

## Acknowledgments

We gratefully acknowledge lab members, Drs. Yen-Cheng Chen and Ashwanth Francis for assistance with GUV imaging, Dr. You Zhang for functional test of IFITM3-iSNAP and advice on the labeling procedure, Ms. Teddy Khan for assistance with molecular cloning, and Mrs. Hui Wu for assistance with cell culture. We thank Henry Cho (Addexbio Technologies) for technical support with INS-1E cells culture and handling and Dr. Zachary Freyberg (University of Pittsburgh) for advice on handling insulin-secreting cells. We also thank Weirong Yuan (University of Maryland) for technical help with peptide synthesis. This work was funded by the NIH R01 grant AI135806 to G.B.M. J.S. and R.D. thank the MaxSynBio consortium, which is jointly funded by the Federal Ministry of Education and Research (BMBF) of Germany and the Max Planck Society (MPG).

## Author Contributions

XG and GBM conceived this study; XG, JS and MM performed the experiments; WL and XL provided reagents; RD advised on bending modulus measurements; XG and GBM wrote the first draft of the manuscript; all authors read and edited the manuscript.

## References

Amatore, C., Arbault, S., Bouret, Y., Guille, M., Lemaitre, F., & Verchier, Y. (2006). Regulation of exocytosis in chromaffin cells by trans-insertion of lysophosphatidylcholine and arachidonic acid into the outer leaflet of the cell membrane. Chembiochem, 7(12), 1998–2003. doi: 10.1002/cbic.200600194

Amini-Bavil-Olyaee, S., Choi, Y. J., Lee, J. H., Shi, M., Huang, I. C., Farzan, M., & Jung, J. U. (2013). The antiviral effector IFITM3 disrupts intracellular cholesterol homeostasis to block viral entry. Cell Host Microbe, 13(4), 452–464. doi: 10.1016/j.chom.2013.03.006

Angelova, M. I., & Dimitrov, D. S. (1986). Liposome Electroformation. Faraday Discussions, 81, 303-+. doi: DOI 10.1039/dc9868100303

Appourchaux, R., Delpeuch, M., Zhong, L., Burlaud-Gaillard, J., Tartour, K., Savidis, G., … Cimarelli, A. (2019). Functional Mapping of Regions Involved in the Negative Imprinting of Virion Particle Infectivity and in Target Cell Protection by Interferon-Induced Transmembrane Protein 3 against HIV-1. J Virol, 93(2). doi: 10.1128/JVI.01716-18

Bailey, C. C., Huang, I. C., Kam, C., & Farzan, M. (2012). Ifitm3 limits the severity of acute influenza in mice. PLoS Pathog, 8(9), e1002909. doi: 10.1371/journal.ppat.1002909

Bailey, C. C., Kondur, H. R., Huang, I. C., & Farzan, M. (2013). Interferon-induced transmembrane protein 3 is a type II transmembrane protein. J Biol Chem, 288(45), 32184–32193. doi: 10.1074/jbc.M113.514356

Bassereau, P., Jin, R., Baumgart, T., Deserno, M., Dimova, R., Frolov, V. A., … Weikl, T. R. (2018). The 2018 biomembrane curvature and remodeling roadmap. J Phys D Appl Phys, 51(34). doi: 10.1088/1361-6463/aacb98

Brass, A. L., Huang, I. C., Benita, Y., John, S. P., Krishnan, M. N., Feeley, E. M., … Elledge, S. J. (2009). The IFITM proteins mediate cellular resistance to influenza A H1N1 virus, West Nile virus, and dengue virus. Cell, 139(7), 1243–1254. doi: 10.1016/j.cell.2009.12.017

Burns, S. M., Vetere, A., Walpita, D., Dancik, V., Khodier, C., Perez, J., … Altshuler, D. (2015). High-throughput luminescent reporter of insulin secretion for discovering regulators of pancreatic Beta-cell function. Cell Metab, 21(1), 126–137. doi: 10.1016/j.cmet.2014.12.010

Chazal, N., & Gerlier, D. (2003). Virus entry, assembly, budding, and membrane rafts. Microbiology and Molecular Biology Reviews, 67(2), 226-+. doi: 10.1128/Mmbr.67.2.226-237.2003

Chernomordik, L. V., Frolov, V. A., Leikina, E., Bronk, P., & Zimmerberg, J. (1998). The pathway of membrane fusion catalyzed by influenza hemagglutinin: restriction of lipids, hemifusion, and lipidic fusion pore formation. J Cell Biol, 140(6), 1369–1382. doi: 10.1083/jcb.140.6.1369

Chernomordik, L. V., & Kozlov, M. M. (2003). Protein-lipid interplay in fusion and fission of biological membranes. Annu Rev Biochem, 72, 175–207.

Chernomordik, L. V., Leikina, E., Frolov, V., Bronk, P., & Zimmerberg, J. (1997). An early stage of membrane fusion mediated by the low pH conformation of influenza hemagglutinin depends upon membrane lipids. J Cell Biol, 136(1), 81–93. doi: 10.1083/jcb.136.1.81

Chernomordik, L. V., Melikyan, G. B., & Chizmadzhev, Y. A. (1987). Biomembrane fusion: a new concept derived from model studies using two interacting planar lipid bilayers. Biochim Biophys Acta, 906(3), 309–352.

Chesarino, N. M., Compton, A. A., McMichael, T. M., Kenney, A. D., Zhang, L., Soewarna, V., … Yount, J. S. (2017). IFITM3 requires an amphipathic helix for antiviral activity. EMBO Rep, 18(10), 1740–1751. doi: 10.15252/embr.201744100

Desai, T. M., Marin, M., Chin, C. R., Savidis, G., Brass, A. L., & Melikyan, G. B. (2014). IFITM3 restricts influenza A virus entry by blocking the formation of fusion pores following virus-endosome hemifusion. PLoS Pathog, 10(4), e1004048. doi: 10.1371/journal.ppat.1004048

Diamond, M. S., & Farzan, M. (2013). The broad-spectrum antiviral functions of IFIT and IFITM proteins. Nat Rev Immunol, 13(1), 46–57. doi: 10.1038/nri3344

Drin, G., & Antonny, B. (2010). Amphipathic helices and membrane curvature. Febs Letters, 584(9), 1840–1847. doi: 10.1016/j.febslet.2009.10.022

Everitt, A. R., Clare, S., McDonald, J. U., Kane, L., Harcourt, K., Ahras, M., … Kellam, P. (2013). Defining the range of pathogens susceptible to Ifitm3 restriction using a knockout mouse model. PLoS One, 8(11), e80723. doi: 10.1371/journal.pone.0080723

Everitt, A. R., Clare, S., Pertel, T., John, S. P., Wash, R. S., Smith, S. E., … Kellam, P. (2012). IFITM3 restricts the morbidity and mortality associated with influenza. Nature, 484(7395), 519–523. doi: 10.1038/nature10921

Feeley, E. M., Sims, J. S., John, S. P., Chin, C. R., Pertel, T., Chen, L. M., … Brass, A. L. (2011). IFITM3 inhibits influenza A virus infection by preventing cytosolic entry. PLoS Pathog, 7(10), e1002337. doi: 10.1371/journal.ppat.1002337

Francis, A. C., Marin, M., Shi, J., Aiken, C., & Melikyan, G. B. (2016). Time-Resolved Imaging of Single HIV-1 Uncoating In Vitro and in Living Cells. PLoS Pathog, 12(6), e1005709. doi: 10.1371/journal.ppat.1005709

Fu, B., Wang, L., Li, S., & Dorf, M. E. (2017). ZMPSTE24 defends against influenza and other pathogenic viruses. J Exp Med, 214(4), 919–929. doi: 10.1084/jem.20161270

Fuller, N., & Rand, R. P. (2001). The influence of lysolipids on the spontaneous curvature and bending elasticity of phospholipid membranes. Biophysical Journal, 81(1), 243–254. doi: Doi 10.1016/S0006-3495(01)75695-0

Girard, P., Pecreaux, J., Lenoir, G., Falson, P., Rigaud, J. L., & Bassereau, P. (2004). A new method for the reconstitution of membrane proteins into giant unilamellar vesicles. (vol 87, pg 419, yr 2004). Biophysical Journal, 87(3), 2098–2098. doi: 10.1529/biophysj.104.900104

Gracia, R. S., Bezlyepkina, N., Knorr, R. L., Lipowsky, R., & Dimova, R. (2010). Effect of cholesterol on the rigidity of saturated and unsaturated membranes: fluctuation and electrodeformation analysis of giant vesicles. Soft Matter, 6(7), 1472–1482. doi: 10.1039/b920629a

Harrison, S. C. (2008). Viral membrane fusion. Nat Struct Mol Biol, 15(7), 690–698. doi: 10.1038/nsmb.1456

Helfrich, W. (1973). Elastic Properties of Lipid Bilayers - Theory and Possible Experiments. Zeitschrift Fur Naturforschung C-a Journal of Biosciences, C 28(11-1), 693–703. doi: DOI 10.1515/znc-1973-11-1209

Huang, I. C., Bailey, C. C., Weyer, J. L., Radoshitzky, S. R., Becker, M. M., Chiang, J. J., … Farzan, M. (2011). Distinct patterns of IFITM-mediated restriction of filoviruses, SARS coronavirus, and influenza A virus. PLoS Pathog, 7(1), e1001258. doi: 10.1371/journal.ppat.1001258

Jia, R., Xu, F. W., Qian, J., Yao, Y. F., Miao, C. H., Zheng, Y. M., … Liang, C. (2014). Identification of an endocytic signal essential for the antiviral action of IFITM3. Cellular Microbiology, 16(7), 1080–1093. doi: 10.1111/cmi.12262

John, S. P., Chin, C. R., Perreira, J. M., Feeley, E. M., Aker, A. M., Savidis, G., … Brass, A. L. (2013). The CD225 domain of IFITM3 is required for both IFITM protein association and inhibition of influenza A virus and dengue virus replication. J Virol, 87(14), 7837–7852. doi: 10.1128/JVI.00481-13

Julkowska, M. M., Rankenberg, J. M., & Testerink, C. (2013). Liposome-binding assays to assess specificity and affinity of phospholipid-protein interactions. Methods Mol Biol, 1009, 261–271. doi: 10.1007/978-1-62703-401-2_24

Kaiser, H. J., Lingwood, D., Levental, I., Sampaio, J. L., Kalvodova, L., Rajendran, L., & Simons, K. (2009). Order of lipid phases in model and plasma membranes. Proceedings of the National Academy of Sciences of the United States of America, 106(39), 16645–16650. doi: 10.1073/pnas.0908987106

Karimi, M., Steinkuhler, J., Roy, D., Dasgupta, R., Lipowsky, R., & Dimova, R. (2018). Asymmetric Ionic Conditions Generate Large Membrane Curvatures. Nano Lett, 18(12), 7816–7821. doi: 10.1021/acs.nanolett.8b03584

Kozlov, M. M. (2007). Determination of lipid spontaneous curvature from X-ray examinations of inverted hexagonal phases. Methods Mol Biol, 400, 355–366. doi: 10.1007/978-1-59745-519-0_24

Kozlov, M. M., Leikin, S. L., Chernomordik, L. V., Markin, V. S., & Chizmadzhev, Y. A. (1989). Stalk mechanism of vesicle fusion. Intermixing of aqueous contents. Eur Biophys J, 17(3), 121–129. doi: 10.1007/BF00254765

Levental, I., Lingwood, D., Grzybek, M., Coskun, U., & Simons, K. (2010). Palmitoylation regulates raft affinity for the majority of integral raft proteins. Proceedings of the National Academy of Sciences of the United States of America, 107(51), 22050–22054. doi: 10.1073/pnas.1016184107

Li, K., Markosyan, R. M., Zheng, Y. M., Golfetto, O., Bungart, B., Li, M., … Liu, S. L. (2013). IFITM proteins restrict viral membrane hemifusion. PLoS Pathog, 9(1), e1003124. doi: 10.1371/journal.ppat.1003124

Lin, T. Y., Chin, C. R., Everitt, A. R., Clare, S., Perreira, J. M., Savidis, G., … Brass, A. L. (2013). Amphotericin B increases influenza A virus infection by preventing IFITM3-mediated restriction. Cell Rep, 5(4), 895–908. doi: 10.1016/j.celrep.2013.10.033

Ling, S., Zhang, C., Wang, W., Cai, X., Yu, L., Wu, F., … Tian, C. (2016). Combined approaches of EPR and NMR illustrate only one transmembrane helix in the human IFITM3. Sci Rep, 6, 24029. doi: 10.1038/srep24029

Lingwood, D., Ries, J., Schwille, P., & Simons, K. (2008). Plasma membranes are poised for activation of raft phase coalescence at physiological temperature. Proceedings of the National Academy of Sciences of the United States of America, 105(29), 10005–10010. doi: 10.1073/pnas.0804374105

Martyna, A., Bahsoun, B., Badham, M. D., Srinivasan, S., Howard, M. J., & Rossman, J. S. (2017). Membrane remodeling by the M2 amphipathic helix drives influenza virus membrane scission. Scientific Reports, 7. doi: ARTN 44695 10.1038/srep44695

McMahon, H. T., & Gallop, J. L. (2005). Membrane curvature and mechanisms of dynamic cell membrane remodelling. Nature, 438(7068), 590–596. doi: 10.1038/nature04396

Merglen, A., Theander, S., Rubi, B., Chaffard, G., Wollheim, C. B., & Maechler, P. (2004). Glucose sensitivity and metabolism-secretion coupling studied during two-year continuous culture in INS-1E insulinoma cells. Endocrinology, 145(2), 667–678. doi: 10.1210/en.2003-1099

Mudhasani, R., Tran, J. P., Retterer, C., Radoshitzky, S. R., Kota, K. P., Altamura, L. A., … Bavari, S. (2013). IFITM-2 and IFITM-3 but not IFITM-1 restrict Rift Valley fever virus. J Virol, 87(15), 8451–8464. doi: 10.1128/JVI.03382-12

Parasassi, T., De Stasio, G., Dubaldo, A., & Gratton, E. (1990). Phase Fluctuation in Phospholipid-Membranes Revealed by Laurdan Fluorescence. Biophysical Journal, 57(6), 1179–1186. doi: Doi 10.1016/S0006-3495(90)82637-0

Rajendran, L., & Simons, K. (2005a). Lipid rafts and membrane dynamics. Journal of Cell Science, 118(6), 1099–1102. doi: 10.1242/jcs.01681

Rajendran, L., & Simons, K. (2005b). Lipid rafts and membrane dynamics. Journal of Cell Science, 118(Pt 6), 1099–1102. doi: 10.1242/jcs.01681

Rigaud, J. L., & Levy, D. (2003). Reconstitution of membrane proteins into liposomes. Liposomes, Pt B, 372, 65-86. Retrieved from <Go to ISI>://WOS:000186734500004

Rossman, J. S., Jing, X. H., Leser, G. P., & Lamb, R. A. (2010). Influenza Virus M2 Protein Mediates ESCRT-Independent Membrane Scission. Cell, 142(6), 902–913. doi: 10.1016/j.cell.2010.08.029

Stachowiak, J. C., Hayden, C. C., & Sasaki, D. Y. (2010). Steric confinement of proteins on lipid membranes can drive curvature and tubulation. Proceedings of the National Academy of Sciences of the United States of America, 107(17), 7781–7786. doi: 10.1073/pnas.0913306107

Steinkuhler, J., Knorr, R. L., Zhao, Z., Bhatia, T., Bartelt, S. M., Wegner, S., … Lipowsky, R. (2020). Controlled division of cell-sized vesicles by low densities of membrane-bound proteins. Nat Commun, 11(1), 905. doi: 10.1038/s41467-020-14696-0

Steinkuhler, J., Sezgin, E., Urbancic, I., Eggeling, C., & Dimova, R. (2019). Mechanical properties of plasma membrane vesicles correlate with lipid order, viscosity and cell density. Communications Biology, 2. doi:UNSP 337 10.1038/s42003-019-0583-3

Stiasny, K., & Heinz, F. X. (2004). Effect of membrane curvature-modifying lipids on membrane fusion by tick-borne encephalitis virus. J Virol, 78(16), 8536–8542. doi: 10.1128/JVI.78.16.8536-8542.2004

Suddala, K. C., Lee, C. C., Meraner, P., Marin, M., Markosyan, R. M., Desai, T. M., … Melikyan, G. B. (2019). Interferon-induced transmembrane protein 3 blocks fusion of sensitive but not resistant viruses by partitioning into virus-carrying endosomes. PLoS Pathog, 15(1), e1007532. doi: 10.1371/journal.ppat.1007532

Tartour, K., Nguyen, X. N., Appourchaux, R., Assil, S., Barateau, V., Bloyet, L. M., … Cimarelli, A. (2017). Interference with the production of infectious viral particles and bimodal inhibition of replication are broadly conserved antiviral properties of IFITMs. PLoS Pathog, 13(9), e1006610. doi: 10.1371/journal.ppat.1006610

Vanblitterswijk, W. J., Vanhoeven, R. P., & Vandermeer, B. W. (1981). Lipid Structural Order Parameters (Reciprocal of Fluidity) in Biomembranes Derived from Steady-State Fluorescence Polarization Measurements. Biochimica Et Biophysica Acta, 644(2), 323–332. doi: Doi 10.1016/0005-2736(81)90390-4

Weinberger, A., Tsai, F. C., Koenderink, G. H., Schmidt, T. F., Itri, R., Meier, W., … Marques, C. (2013). Gel-assisted formation of giant unilamellar vesicles. Biophysical Journal, 105(1), 154–164. doi: 10.1016/j.bpj.2013.05.024

Wesolowska, O., Michalak, K., Maniewska, J., & Hendrich, A. B. (2009). Giant unilamellar vesicles - a perfect tool to visualize phase separation and lipid rafts in model systems. Acta Biochimica Polonica, 56(1), 33-39. Retrieved from <Go to ISI>://WOS:000264798400004

Wrensch, F., Winkler, M., & Pohlmann, S. (2014). IFITM Proteins Inhibit Entry Driven by the MERS-Coronavirus Spike Protein: Evidence for Cholesterol-Independent Mechanisms. Viruses, 6(9), 3683–3698. doi: 10.3390/v6093683

Wu, X., Spence, J. S., Das, T., Yuan, X., Chen, C., Zhang, Y., … Peng, T. (2020). Site-Specific Photo-Crosslinking Proteomics Reveal Regulation of IFITM3 Trafficking and Turnover by VCP/p97 ATPase. Cell Chem Biol. doi: 10.1016/j.chembiol.2020.03.004

Yang, S. T., Kreutzberger, A. J. B., Kiessling, V., Ganser-Pornillos, B. K., White, J. M., & Tamm, L. K. (2017). HIV virions sense plasma membrane heterogeneity for cell entry. Science Advances, 3(6). doi: ARTN e1700338 10.1126/sciadv.1700338

Yount, J. S., Moltedo, B., Yang, Y. Y., Charron, G., Moran, T. M., Lopez, C. B., & Hang, H. C. (2010). Palmitoylome profiling reveals S-palmitoylation-dependent antiviral activity of IFITM3. Nature Chemical Biology, 6(8), 610–614. doi: 10.1038/Nchembio.405

Zhang, Y. H., Zhao, Y., Li, N., Peng, Y. C., Giannoulatou, E., Jin, R. H., … Dong, T. (2013). Interferon-induced transmembrane protein-3 genetic variant rs12252-C is associated with severe influenza in Chinese individuals. Nat Commun, 4, 1418. doi: 10.1038/ncomms2433

Zhang, Z., Liu, J., Li, M., Yang, H., & Zhang, C. (2012). Evolutionary dynamics of the interferon-induced transmembrane gene family in vertebrates. PLoS One, 7(11), e49265. doi: 10.1371/journal.pone.0049265

