## Supplemental Figures and Legends for "Interferon-Induced Transmembrane Protein 3 Blocks Fusion of Diverse Enveloped Viruses by Locally Altering Mechanical Properties of Cell Membranes"

### Supplementary Figures and Legends

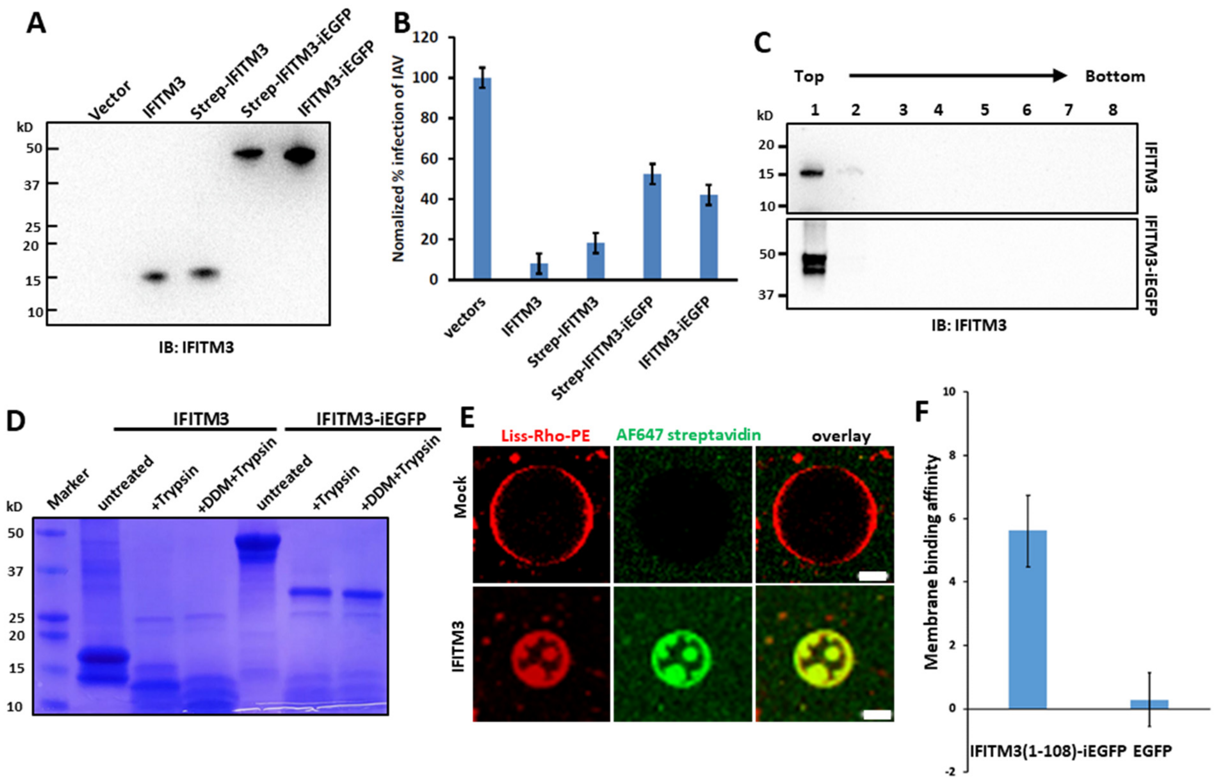


**Figure S1. IFITM3 induces negative membrane curvature *in vitro*.** (A) Expression levels of different IFITM3 constructs in transfected HEK 293T/17. HEK 293T/17 cells transfected with an empty Vector, unlabeled IFITM3, Strep-tagged IFITM3, IFITM3-iEGFP, or Strep-tagged IFITM3-iEGFP were lysed and analyzed by Western blotting using a rabbit antibody against the N-terminal region of IFITM3. (B) IAV pseudovirus infection is inhibited in HEK 293T/17 cells expressing IFITM3 constructs described in (A). The extent of infection by IAV pseudovirus carrying the luciferase gene was measured by luciferase activity is shown as a percentage of infection in cells transfected with empty vector. Data represent mean ± SD (n = 3). (C) IFITM3- and IFITM3-iEGFP-reconstituted LUVs were subjected to floatation analysis. After density gradient centrifugation, collected fractions were analyzed by SDS–PAGE and Western blotting using rabbit anti-IFITM3 antibody. (D) IFITM3- and IFITM3-iEGFP-reconstituted LUVs were incubated with trypsin at 37°C for 30 min. As a control, 0.2% n-Dodecyl-B-D-maltoside (DDM) was added with trypsin to permeabilize LUV membrane. The samples were analyzed by SDS-PAGE and stained with Coomassie blue. (E) Representative GUVs prepared from mock-treated LUVs (without protein, top) or from Strep-tag IFITM3-reconstituted LUVs (bottom) were stained with Streptavidin AlexaFluor-647. Scale bars 10 μm. (F) Membrane binding affinity of IFITM3(1-108)-iEGFP. Two mM of LUVs (99.0 mol % POPC, 0.5 mol % cholesterol, 0.5 mol % Liss-Rho-PE) were incubated at room temperature with 1 μM IFITM3(1-108)-iEGFP or EGFP for 10 min and then spun down by ultra-centrifugation. The binding affinity of protein was deduced from the reduction of EGFP fluorescence in supernatant and normalized by comparing with the fluorescence of input samples. Data represent mean ± SD of the results of three independent experiments.

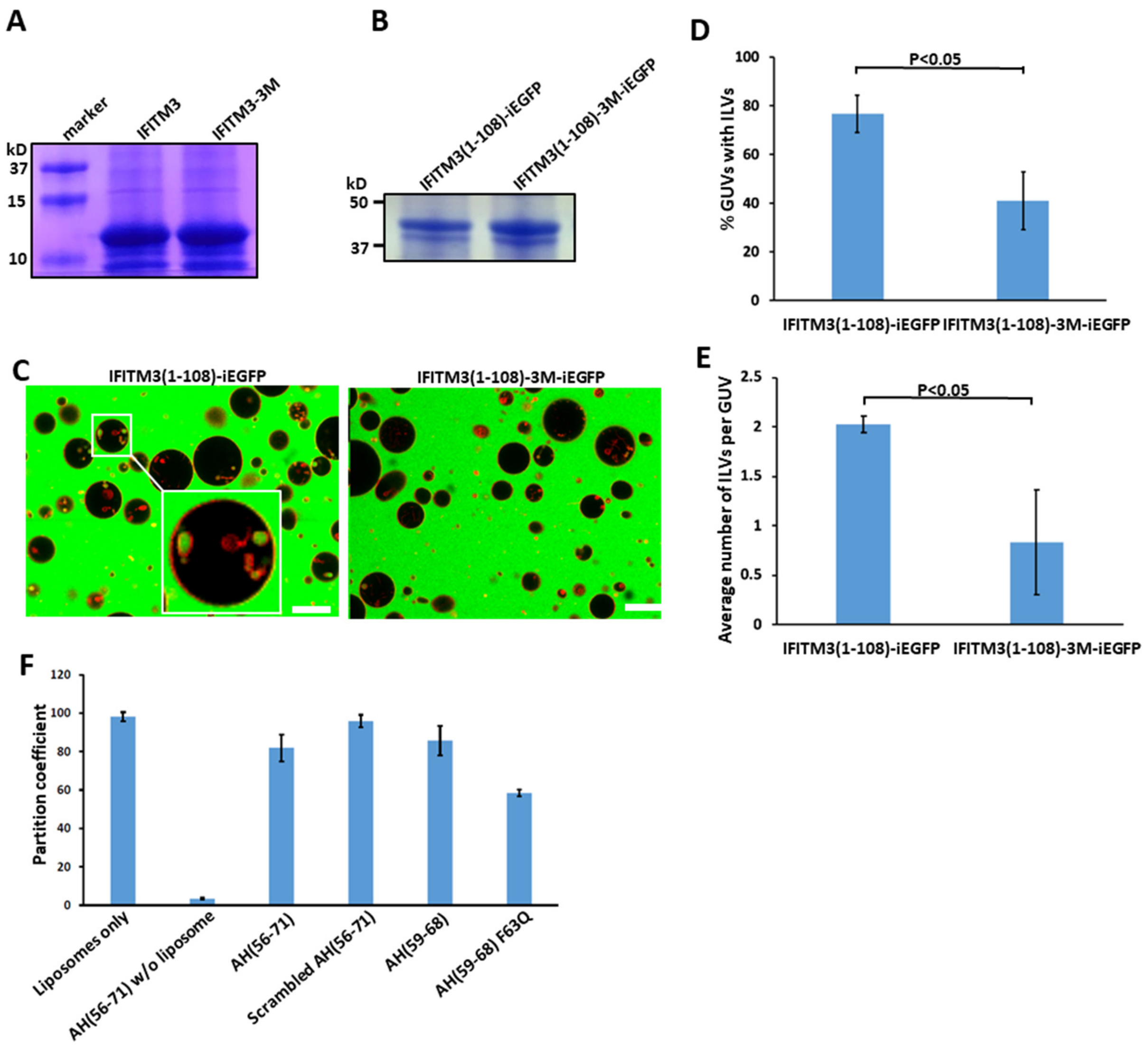


**Figure S2. IFITM3 amphipathic helix is responsible for induction of negative membrane curvature.** (A) Coomassie blue staining of purified IFITM3 and IFITM3-3M mutant. (B) Coomassie blue staining of purified IFITM3(1-108)-iEGFP and IFITM3(1-108)-3M-iEGFP. (C) GUVs were incubated with 20 µM IFITM3(1-108)-iEGFP or IFITM3(1-108)-3M-iEGFP for 30 min and imaged. Scale bars 10 µm. (D) Quantification of inward budding showing the percentage of GUVs, prepared and treated as in (C), with at least one intraluminal vesicle (ILV) containing EGFP. Data represent mean ± SD of the results of three independent experiments, with 13 GUVs analyzed per sample in each experiment. (E) As in (D), but the plots represent the average number of ILVs per GUV. (F) Partition coefficients of peptides. Two mM of LUVs (99.0 mol % POPC, 0.5 mol % cholesterol, 0.5 mol % Liss-Rho-PE) were incubated at room temperature with 40 µM of Cy5-labeled peptides for 10 min and spun down by ultra-centrifugation. The partition coefficient of peptides was deduced from reduction of Cy5 fluorescence in supernatant and normalized to the fluorescence input. Data represent mean ± SD of the results of two independent experiments.

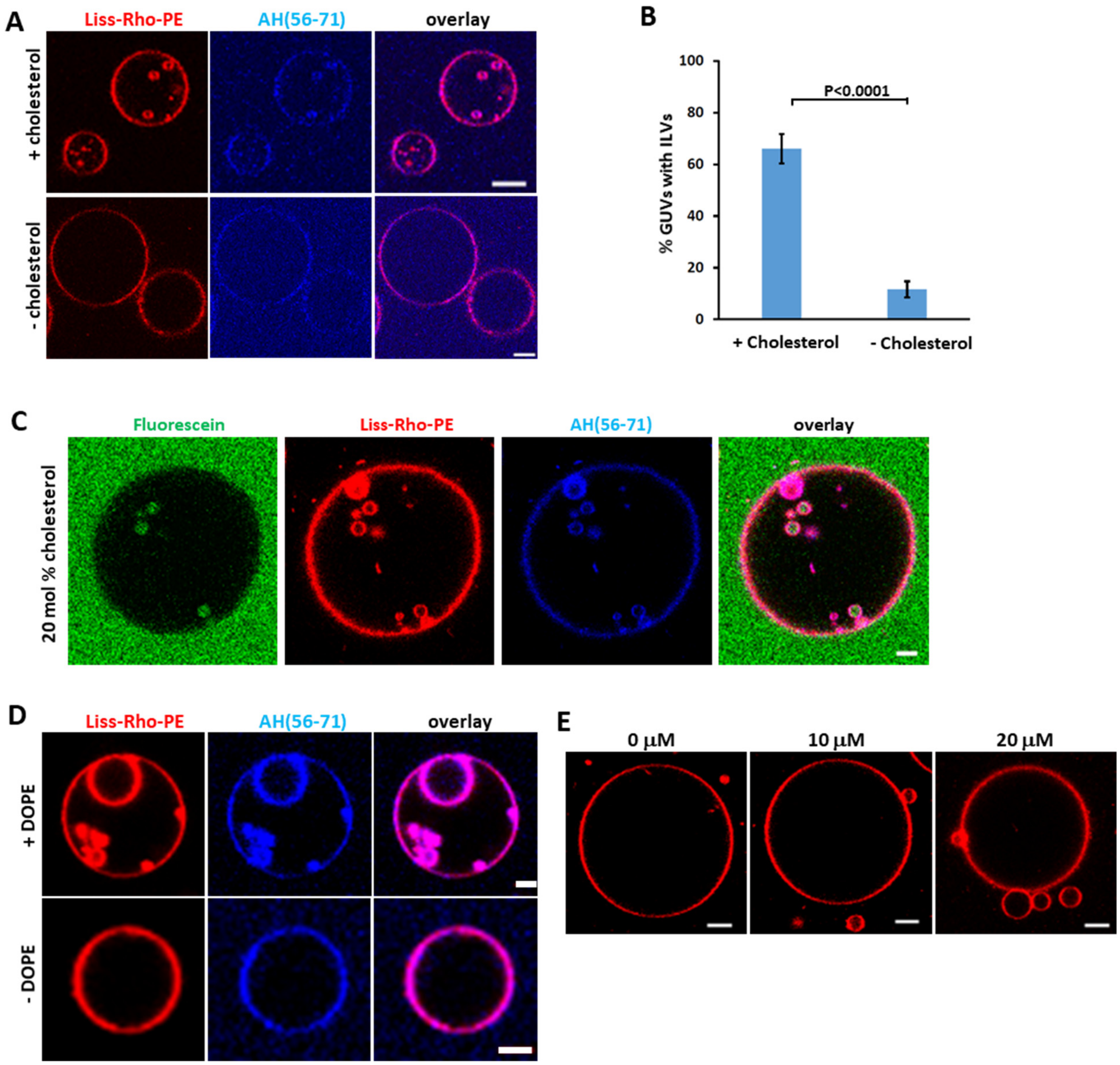


**Figure S3. Negative membrane curvature induced by IFITM3 is facilitated by cholesterol, POPE and counteracted by lyso-lipids.** (A) GUVs with cholesterol (99.0 mol % POPC, 0.5 mol % cholesterol, 0.5 mol % Liss-Rho-PE) or GUVs without cholesterol (99.5 mol % POPC, 0.5 mol % Liss-Rho-PE) were incubated with 10 μM AH(56-71) for 30 min and imaged. Scale bars 10 μm. (B) Quantification of inward budding showing the percentage of GUVs, prepared and treated as in (A), with at least one intraluminal vesicle (ILV) containing Cy5-labeled peptide. Data represent mean ± SD of three independent experiments, with at least 45 GUVs analyzed per sample in each experiment. (C) GUVs (79.5 mol % POPC, 20 mol % cholesterol, 0.5 mol % Liss-Rho-PE) were incubated with 10 μM AH(56-71) for 30 min and imaged. Scale bars 5 μm. (D) GUVs without POPE (99.0 mol % POPC, 0.5 mol % cholesterol, 0.5 mol % Liss-Rho-PE) or with POPE (79.5 mol % POPC, 20 mol % DOPE, 0.5 mol % Liss-Rho-PE) were incubated with 10 μM AH(56-71) for 30 min and imaged. Scale bars 5 μm. (E) GUVs (99.0 mol % POPC, 0.5 mol % cholesterol, 0.5 mol % Liss-Rho-PE) were treated with an indicated concentration of LPC (+LPC) for 30 min and imaged. Scale bars 10 μm.

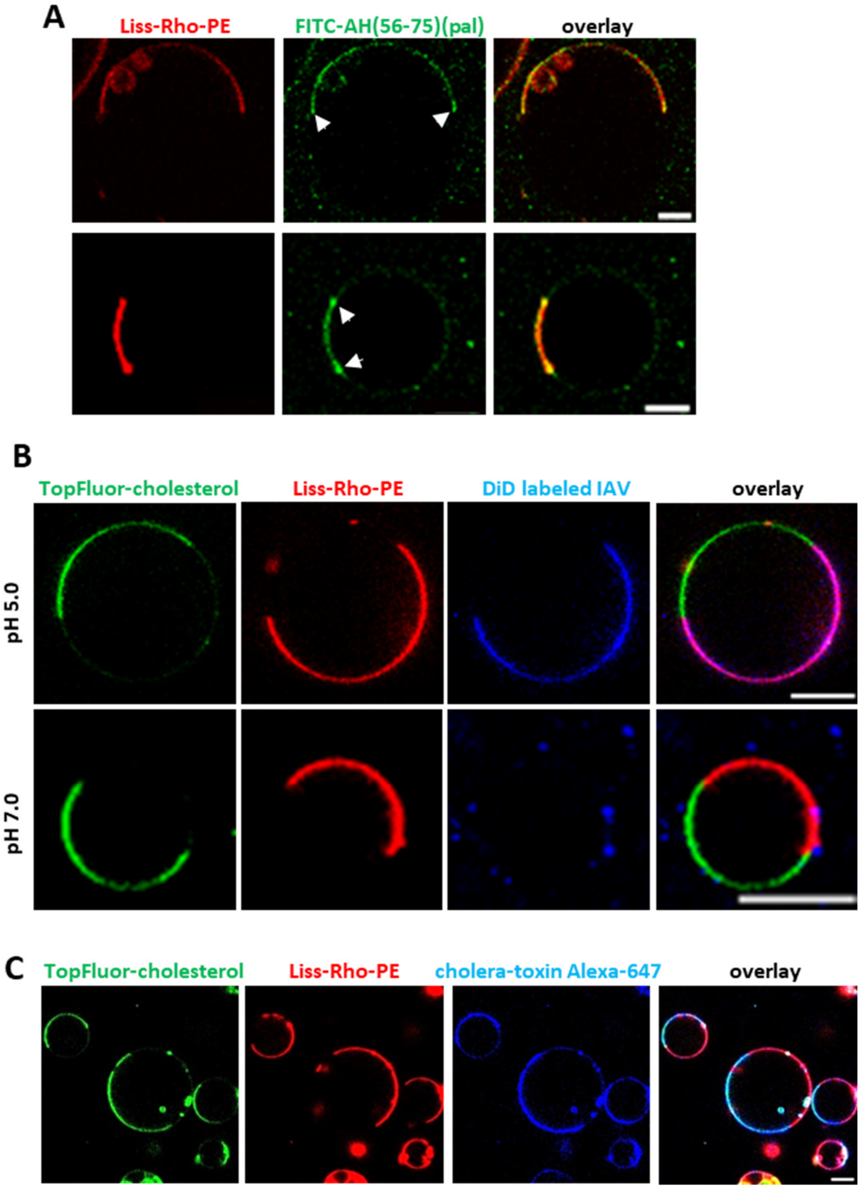


**Figure S4. IFITM3 partitions into liquid-disordered membrane domains that support IAV fusion.** (A) Shown are two examples of phase-separated GUVs (33.3 mol % DOPC, 33.3 mol % SM, 32.4 mol % cholesterol, 0.5 mol % TopFluor-cholesterol and 0.5% Liss-Rho-PE) incubated with 10 µM of FITC-labeled palmitoylated AH (FITC-AH(56-75)(pal)) for 30 min. Arrows indicate accumulation of FITC-AH(56-75)(pal) peptide at the phase boundary. Scale bars 5 µm. (B) Phase-separated GUVs (33.3 mol % DOPC, 33.3 mol % SM, 30.4 mol % cholesterol, 2% GM1, 0.5 mol % TopFluor-cholesterol and 0.5% Liss-Rho-PE) were mixed with DiD-labeled IAV. Lipids mixing between IAV and GUV was triggered by addition of a predetermined amount of citrate buffer to achieve the final pH of 5.0 and samples were immediately imaged. PBS (pH 7.2) was used as control. Scale bars 10 µm. (C) GM1 in phase-separated GUVs (33.3 mol % DOPC, 33.3 mol % SM, 30.4 mol % cholesterol, 2% GM1, 0.5 mol % TopFluor-cholesterol and 0.5% Liss-Rho-PE) was stained with cholera-toxin B Alexa-647. Scale bars 10 µm.

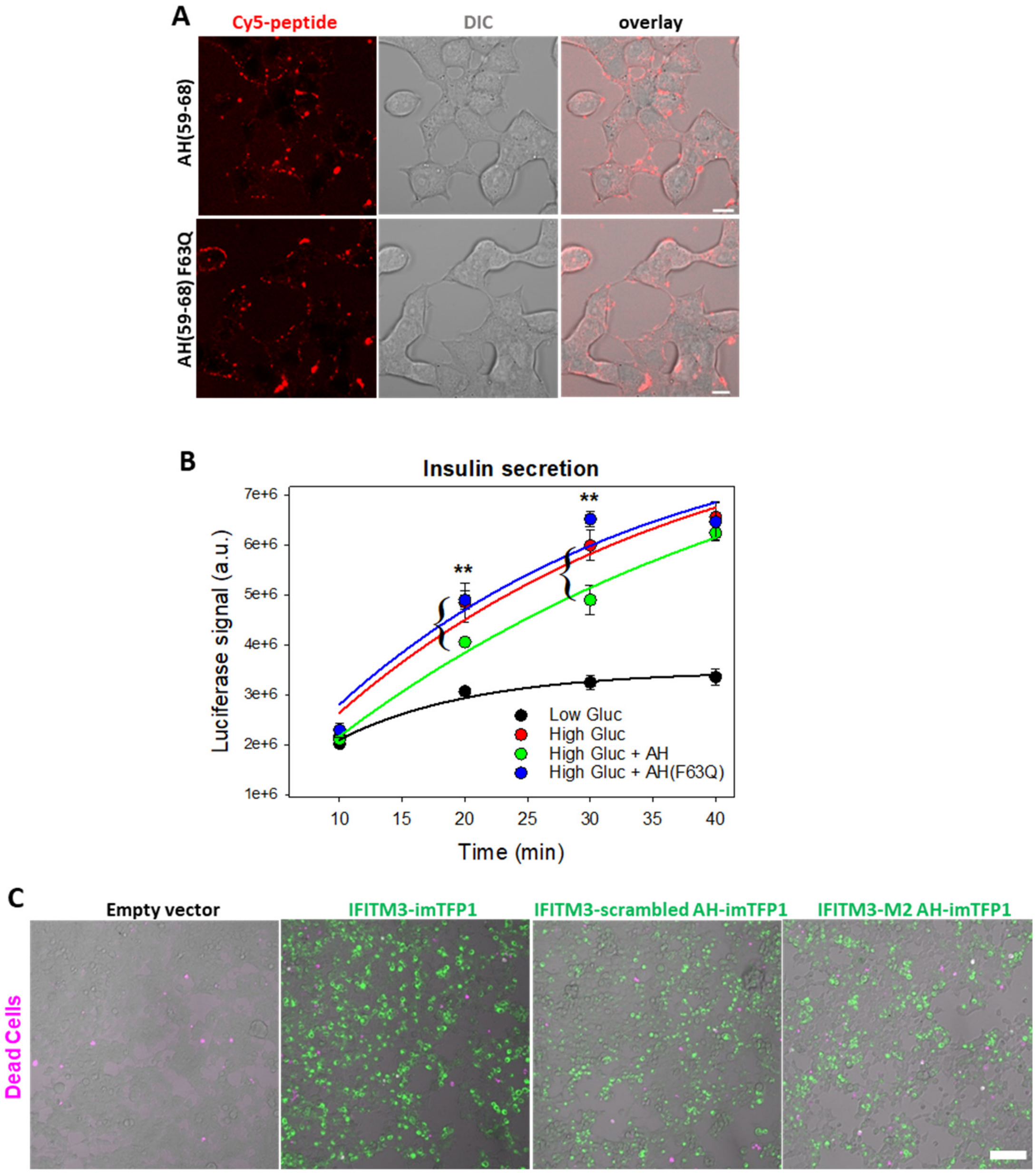


**Figure S5. IFITM3 AH is sufficient to inhibit membrane fusion.** (A) INS-1E cells were treated with 5 μM AH(59-68) or 10 μM AH(59-68) F63Q for 5 min and imaged. Scale bars 10 μm. (B) Time course of glucose-stimulated insulin secretion in INS-1E cells transduced with proinsulin-luciferase fusion construct. INS-1E cells were preincubated for 5 min with 5 μM AH(59-68) or 10 μM AH(59-68) F63Q and stimulated for 20 min with a high glucose (20 mM, HG) buffer or low glucose (2.8 mM, LG) as control. Luciferase activity was measured for each time point by adding the coelenterazine substrate to the supernatant and reading on a Luminescence counter. Data represent mean ± SD of the results of two independent experiments. **, p<0.01 for HG and HG + AH(59-68) samples determined by the Student's test. (C) HEK 293T/17 cells expressing wild-type or scrambled imTFP1tagged IFITM3 or IFITM3 M2 AH chimera were infected by IAV and cell viability was tested using a LIVE/DEAD™ Fixable Far Red Dead Cell Stain Kit 12 h after infection. Dead cells are shown in magenta. Scale bar 100 μm.
